# Multiple ShKT domain-containing MUL-1 proteins act as redox-responsive modulators of oxidative stress signaling in *C. elegans*

**DOI:** 10.64898/2026.05.03.722560

**Authors:** Emilio Carranza-Garcia, Abe Gayle Santos, Kyoung-Hye Yoon, Anton Gartner

**Affiliations:** Center for Genomic Integrity, Institute for Basic Science, UNIST-gil 50, Ulsan 44919, Republic of Korea; Department of Physiology, Yonsei University Wonju College of Medicine, 20 Ilsan-ro, Wonju, South Korea; Organelle Medicine Research Center, Yonsei University Wonju College of Medicine, 20 Ilsan-ro, Wonju, South Korea; Department of Global Medical Science, Yonsei University Wonju College of Medicine, 20 Ilsan-ro, Wonju, South Korea; Department for Health Science and Technology, Ulsan National Institute of Science and Technology (UNIST), UNIST-gil 50, Ulsan 44919, Republic of Korea

**Keywords:** *C. elegans*, oxidative stress, ionizing irradiation, MUL-1, ShKT domain, stress signaling

## Abstract

Organismal survival depends on coordinated responses to oxidative stress and DNA damage. Using *Caenorhabditis elegans*, we investigate *mul-1,* a robust transcriptional target of ionizing radiation and reactive oxygen species. Although annotated as a mucin, MUL-1 is a small ShKT domain-containing protein belonging to an invertebrate expanded family of cysteine-rich proteins. *mul-1* is selectively induced by oxidative stress, including IR, hydrogen peroxide (H_2_O_2_), *Pseudomonas aeruginosa* infection, or loss of the peroxiredoxin PRDX-2, via the p38 MAPK-ATF-7 pathway in intestinal cells. Loss of *mul-1* and its paralogs increases ROS accumulation, oxidative stress sensitivity, and CEP-1/p53 dependent germ cell apoptosis. Combined deletion of *mul-1* paralogs causes constitutive apoptosis, reduced fecundity, and compensatory activation of DAF-16/Foxo and SKN-1/Nrf2 stress response pathways. Together with genetic analysis of SYSM-1, these findings suggest MUL-1-like ShKT proteins buffer oxidative stress.

## Introduction

Organismal survival depends on the activation of coordinated stress response pathways. Ionizing radiation (IR) and reactive oxygen species (ROS) are among the most potent inducers of cellular stress, triggering DNA damage and oxidative insults. The nematode *Caenorhabditis elegans* provides a genetically tractable model to dissect the regulation and functional impact of these pathways at the organismal level. Although IR can directly induce DNA strand breaks, most DNA damage associated with IR exposure arises indirectly through ROS generation. ROS include superoxide (O_2_^-^), hydrogen peroxide (H_2_O_2_), and hydroxyl radicals (•OH), produced when radiation interacts with cellular water and organic molecules (Roots & Okada, 1975; Ward, 1994). In addition to oxidizing bases, generating abasic sites, and causing DNA strand breaks, ROS also inflict cellular damage by oxidizing metabolites, lipids, and proteins (Dalle-Donne et al., 2006; Yohe & Davies, 2014). ROS are also generated endogenously, for instance, through mitochondrial electron leakage and NADPH oxidase activity (Forman et al., 2010). Among ROS, H_2_O_2_ is a precursor to highly reactive hydroxyl radicals generated via Fenton chemistry (Forman & Zhang, 2021; Koppenol, 1993), but can also act as a signaling molecule (Forman et al., 2010; Miranda-Vizuete & Veal, 2017). For instance, in *C. elegans,* H_2_O_2_ regulates FLP-1 neuropeptide release from AIY interneurons during diet-induced stress response in the gut (Jia & Sieburth, 2021), and this response is potentiated by the H_2_O_2_-dependent release of the FLP-2 peptide from the intestine (Jia et al., 2024). Elevated ROS in AWC neurons causes NLP-1 peptide secretion, which induces the mitochondrial unfolded protein response in the gut and reduces its digestive capacity (Liu et al., 2024). Cellular detoxification of H_2_O_2_ is primarily mediated by antioxidant enzymes such as superoxide dismutases, peroxiredoxins, and glutathione peroxidases, which rely on conserved cysteine residues or thiol-containing cofactors for redox cycling (Aranda-Rivera et al., 2022; Juan et al., 2021).

Transcriptomic analyses following IR in *C. elegans* revealed no induction of canonical DNA repair genes (Greiss et al., 2008). Among the DNA damage response genes, only the pro-apoptotic BH3-only genes *egl-1* and *ced-13*, both CEP-1/p53 targets, and required for DNA damage-induced germ cell apoptosis, were upregulated (Greiss et al., 2008; Schumacher et al., 2005). In contrast, a broad CEP-1/p53 independent transcriptional activation of oxidative stress-related and innate immunity-associated genes was observed, many of which are nematode-specific (Greiss et al., 2008). Notably, *mul-1* emerged as the most robustly IR-induced transcript, and its induction requires the conserved p38 MAPK pathway (Greiss et al., 2008; Kimura et al., 2012). Recently, a MUL-1 high-copy transgene was shown to be expressed in the gut, and *mul-1* deletion was associated with reduced sensitivity to *Pseudomonas aeruginosa* infection, possibly by limiting bacterial association with gut epithelium (Hoffman et al., 2020). However, while annotated as a mucin-like gene, MUL-1 lacks some hallmark features of vertebrate mucins, which are typically thousands of amino acids long, highly enriched in serine and threonine, and heavily glycosylated to form gel-like protective barriers in gut epithelia (Johansson et al., 2013). Instead, MUL-1 is a small 259-amino-acid protein composed mainly of five ∼36-42 amino acid ShKT domains (InterPro Entry IPR003582: ShKT domain). Only a 42-amino-acid unstructured region between the two C-terminal ShKT domains is highly enriched in serine/threonine residues. ShKT domains were initially characterized as potent toxins derived from sea anemones that inhibit mammalian potassium channels (Castañeda et al., 1995; Gerdol et al., 2019; Harvey & Vita, n.d.; Shafee et al., 2019; Tudor et al., 1998). Except for the human metalloprotease MMP23, which contains a single ShKT module, this motif is otherwise absent from vertebrate proteomes (Rangaraju et al., 2010). Structurally, ShKT domains are defined by six conserved cysteine residues that form three disulfide bonds, stabilizing a compact two-α-helix fold commonly used to engage and modulate potassium channels (Castañeda et al., 1995; Shafee et al., 2019; Tudor et al., 1998). Given this distinctive organization, we posit that MUL-1 may perform roles unrelated to, or in addition to, those of traditional mucins.

The identification of *mul-1* as an IR-responsive gene is reminiscent of *sysm-1*, a small protein also induced by IR and composed of two ShKT domains (Soltanmohammadi et al., 2022). Like *mul-1*, *sysm-1* induction depends on the p38 MAPK pathway (Soltanmohammadi et al., 2022). Functional studies have shown that SYSM-1 is secreted from the intestine and is required for germ cell apoptosis following IR, acting in parallel to the *C. elegans* CEP-1/p53 pathway. Notably, the induction of the two pro-apoptotic BH3 domain-only genes, *egl-1* and *ced-13* remains intact in *sysm-1* mutants, suggesting that SYSM-1 conveys stress signals across tissues, independent of CEP-1 transcriptional activity (Soltanmohammadi et al., 2022).

Here, we employed a *mul-1* transcriptional reporter as an inroad to dissect regulatory circuits involved in the oxidative stress response. *mul-1* is induced by IR, H_2_O_2_, *Pseudomonas* infection and loss of the peroxiredoxin PRDX-2. Peroxiredoxins are abundant cysteine-based peroxide reductases that detoxify H_2_O_2_ through the oxidation of conserved N-terminal cysteines to sulfenic acid, followed by disulfide bond formation with a receiving cysteine (Rhee, 2016). We found that *mul-1* expression is induced in the intestine and depends on p38 signaling and its downstream transcription factor, ATF-7. *mul-1* mutants are hypersensitive to oxidative stress and exhibit increased p53-dependent germ cell apoptosis upon IR. MUL-1 belongs to a family of proteins expanded in invertebrates, and we included the three most closely related paralogs, as well as related *sysm-1,* in our analysis. MUL-1 family quadruple mutants display a further increase in radiation-induced apoptosis, and excessive apoptosis occurs even in the absence of IR. Furthermore, both *mul-1* and the quadruple mutant bypass the apoptosis defect of *sysm-1*. In compound *mul-1* paralog mutants and *prdx-2* single mutants, compensatory DAF-16-dependent SOD-3 and SKN-1-dependent GST-4 induction occurs even in the absence of exogenous stress. We argue that MUL-1-like proteins are part of a regulatory circuit that have a key role in the organismal responses to oxidative stress. We hypothesize that MUL1-like genes may act via their ShKT domains as scavengers or rheostats of oxidative damage.

## Results

### Transcriptional regulation of *mul-1*

To assess if and where *mul-1* (F49F1.6) is induced upon IR, we developed a transcriptional reporter strain, *mul-1(syb3342)*, in which the coding sequence of *mul-1* is replaced with mCherry fused to histone H2B for fluorescent detection and nuclear targeting. Under control conditions, the basal expression of *mul-1* is predominantly localized to the nuclei of gut cells, with the strongest expression observed in the two anterior-most gut nuclei (Fig. 1A, C), with expression gradually increasing during larval development, reaching a maximum in L4 larvae and adults (Fig. 1A, C-D, black lines). To examine the induction of *mul-1* under DNA-damaging conditions, we exposed *mul-1(syb3342)* animals at all larval stages to 100 Gy of IR and analyzed transcriptional activation 6 hours post-treatment. IR exposure results in a strong induction of *mul-1* across all gut cells at every developmental stage, most notably in the anterior two nuclei (Fig. 1B, C-D, red lines). The induction is dose- and time-dependent, becoming detectable after 2 hours and peaking at 6 hours (Figs. S1, S2). For Western blotting, we generated a knock-in strain with a C-terminal 3xHA tag at the endogenous *mul-1* locus. In untreated controls, MUL-1::3xHA protein was undetectable by immunoblotting, consistent with very low basal expression (Fig. 1). However, 6 hours after IR treatment, a specific ∼35 kDa band corresponding to the predicted molecular weight appears (Fig. S3).

**Figure 1.**
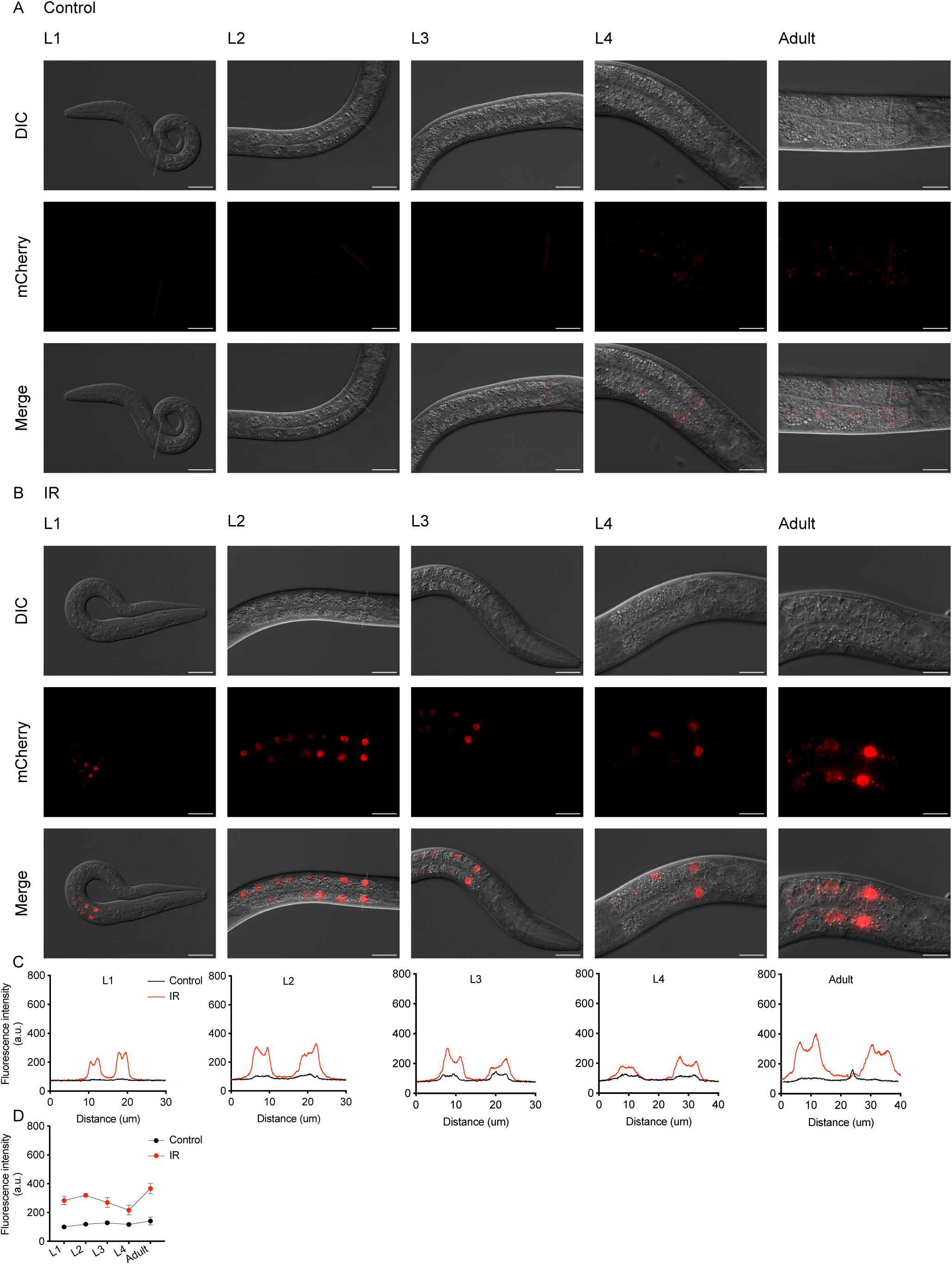
IR induces transcriptional activation of *mul-1* in the intestine of *C. elegans*. (A) Under control conditions, *mul-1* expression in *mul-1(syb3342) IV* animals is detected predominantly in intestinal nuclei, with the strongest signal in the two anterior-most nuclei and a gradual increase during larval development. (B) Six hours after exposure to 100 Gy IR, *mul-1* expression is robustly induced in intestinal nuclei across all larval stages, initiating in the anterior cells and subsequently expanding throughout the gut. (C) Representative fluorescence intensity profiles measured along a transverse line across the first pair of anterior intestinal nuclei in *mul-1(syb3342) IV* animals. (D) Quantification of nuclear fluorescence intensity derived from the maximum intensity peaks corresponding to each nucleus, following optimized line placement to minimize background contributions from adjacent intestinal cells in different focal planes. Scale bars, 20 μm. Statistical analysis was performed using two-way ANOVA with Šidak’s multiple comparisons test. Quantification includes animals from at least three independent experiments (n = 20-30 animals per developmental stage and condition).

To determine if the induction of *mul-1* is specific to IR-induced DNA damage, we tested other genotoxic agents. Neither cisplatin, a DNA crosslinking agent, nor methyl methanesulfonate (MMS), an alkylating agent, induced *mul-1* expression (Fig. 2A-D, F). In contrast, some induction was observed following UV treatment (Fig. 2E, F). UV exposure not only generates cyclobutane pyrimidine dimers (CPDs) and 6-4 photoproducts (6-4 PPs) but also produces ROS through photochemical reactions (Yoshiyama et al., 2023), suggesting that *mul-1* induction might be linked to oxidative stress rather than DNA damage itself. Additionally, no induction was observed after starvation, heat shock, or osmotic stress (Fig. 3A-D, K). However, a strong induction occurred after exposure to H_2_O_2_, a potent ROS generator that produces hydroxyl radicals (·OH) and superoxide anions (O2-) (Kumsta et al., 2011) (Fig. 3E, K). ROS are also produced during normal metabolism, and the two 2-Cys peroxiredoxins, PRDX-2 and PRDX-3, serve as key antioxidants, with PRDX-2 playing a vital role in detoxifying H_2_O_2_. Loss of PRDX-2 results in increased sensitivity to H_2_O_2_, a shortened lifespan, and developmental abnormalities (Kumsta et al., 2011; Oláhová et al., 2008). To investigate whether impaired H_2_O_2_ detoxification can trigger *mul-1* induction, we used CRISPR/Cas9 to introduce the *prdx-2(gk169)* mutation into *mul-1(syb3342)* animals. Under normal conditions, basal *mul-1* expression in *prdx-2(gk169); mul-1(syb3342)* animals was similar to that of controls during early larval stages (Fig. 3F). However, as development progressed, *mul-1* transcription first appeared in anterior gut cells. It gradually expanded along the intestine, eventually resulting in widespread, strong expression in adult animals (Fig. 3G-J, L). Our results suggest that both exogenous and endogenous ROS induce *mul-1* expression.

**Figure 2.**
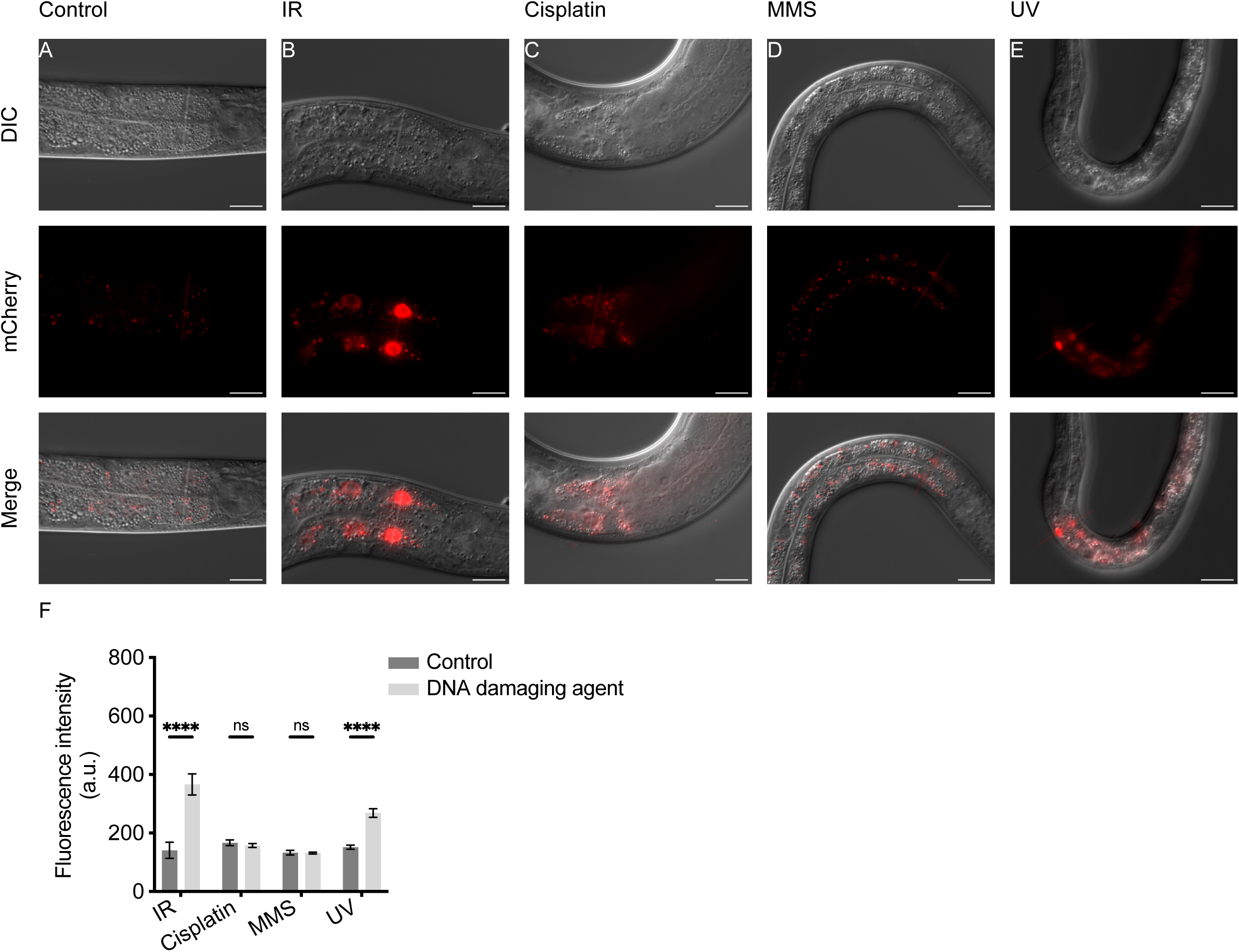
mul-1 expression is selectively induced by IR but not by other DNA-damaging agents. Representative images of *mul-1(syb3342) IV* animals under control (A) conditions or following exposure to IR (B), cisplatin (C), MMS (D), or UV irradiation (E). Robust induction of *mul-1* expression in intestinal nuclei is observed specifically after IR (B) whereas other DNA-damaging agents elicit little or no reporter activation (C-E). Animals were treated with the respective agents at the L4 stage and assayed after 6h after IR (B), 24h after cisplatin treatment (C), 16 hours after MMS treatment (D) and 24hours after UV treatment (E) (Materials and Methods). (F) quantification of nuclear mCherry fluorescence intensity in intestinal cells under the indicated conditions. Scale bars, 20 µm. Data are shown as mean ± SEM and include animals from at least three independent experiments (n = 20-30 animals per condition). Statistical analysis was performed using two-way ANOVA Tukey’s multiple comparisons test.

**Figure 3.**
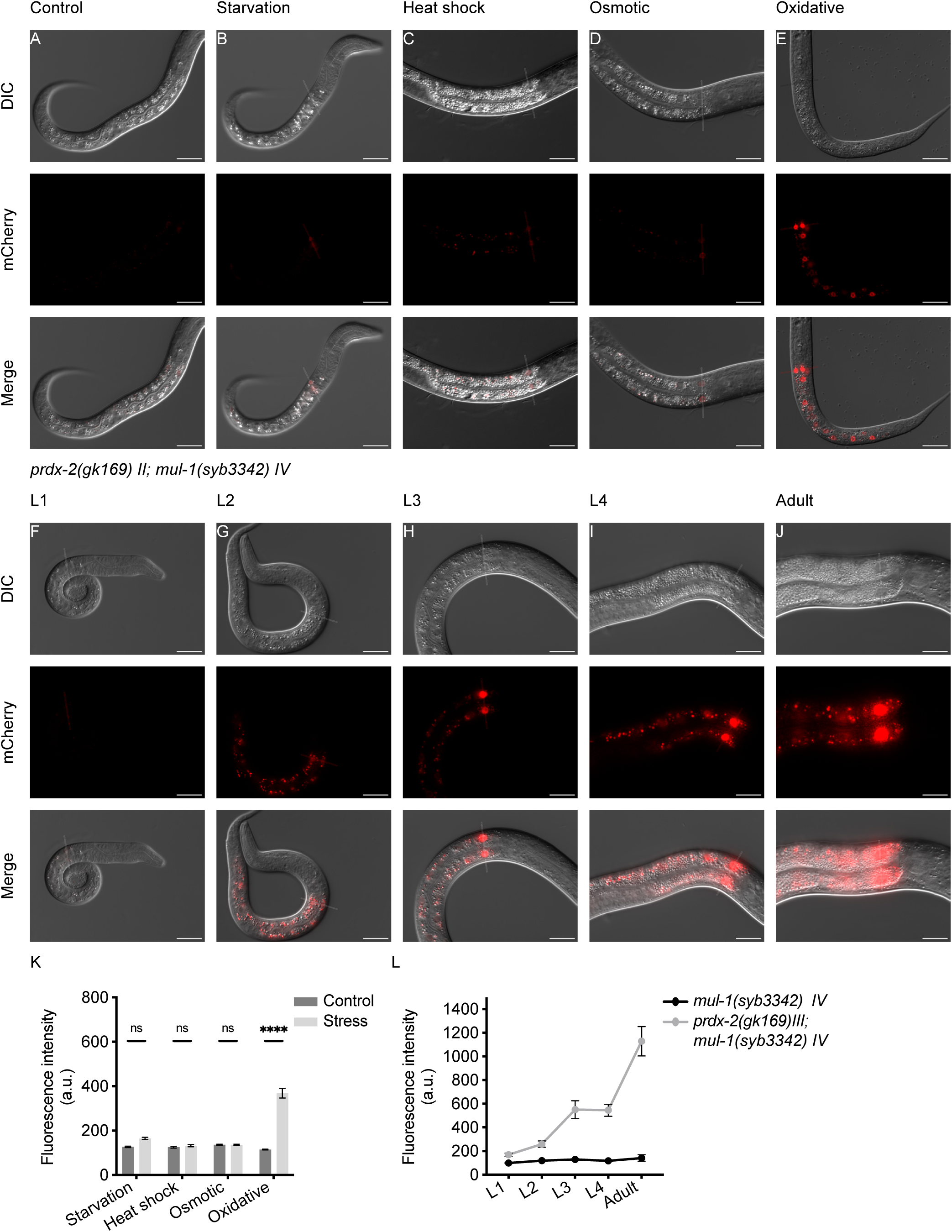
Oxidative stress, but not general stressors, triggers *mul-1* induction. (A-D) Representative images of *mul-1(syb3342) IV* animals subjected to starvation, heat shock, or osmotic stress show no detectable reporter activation in intestinal nuclei. (E) Treatment with H_2_O_2_ induces robust *mul-1* expression in intestinal nuclei. (A-E) All animals were treated at the L1 stage and imaged after 24hours (note that starved and H_2_O_2_ treated worms are developmentally arrested. (F-G) Basal *mul-1* expression in *prdx-2(gk169) II; mul-1(syb3342) IV* animals is comparable to controls during early larval stages (L1-L2). (H-J) As development progresses, *mul-1* activation in *prdx-2* mutants initiates in anterior intestinal nuclei and gradually extends toward posterior regions of the gut. (K) Quantification of fluorescence intensity in anterior intestinal nuclei confirms selective *mul-1* induction by oxidative stress. (L) Comparative quantification under control conditions reveals elevated basal *mul-1* activation in *prdx-2* mutants. Scale bars, 20 µm. Data are shown as mean ± SEM and include animals from at least three independent experiments (n = 20-30 animals per condition). Statistical analysis was performed using two-way ANOVA followed by Tukey’s multiple comparisons test.

### The p38 MAPK pathway and its downstream effector ATF-7 regulate *mul-1*

Next, we examined the role of key stress response pathways in regulating *mul-1*. Previous RNAi studies and quantitative PCR showed that both the p38/PMK-1 and insulin/IGF-1 signaling pathways are necessary for *mul-1* induction after IR treatment (Kimura et al., 2012). Using our *mul-1* transcriptional reporter, we systematically dissected the contribution of the p38/PMK-1 pathway and its downstream effectors, including SKN-1 and ATF-7, as well as the upstream regulator SEK-1, in response to IR-induced DNA damage. In *C. elegans*, the TIR-1-NSY-1-SEK-1-PMK-1 signaling cascade mediates the innate immune response in the gut (Inoue et al., 2005). SEK-1 (stress-activated protein kinase-1), a mitogen-activated protein kinase kinase (MAPKK), acts upstream of p38/PMK-1, modulating its activity through phosphorylation in response to various stress stimuli, including infection, oxidative stress, or environmental insults (Kim et al., 2002). We found that IR-induced *mul-1* upregulation is compromised in *sek-1(km4)* and *pmk-1 (km25)* mutants (Fig. 4A-C, F). We also found that ATF-7, but not the SKN-1 downstream effector, is required for *mul-1* induction (Fig. 4D-F). ATF-7 and SKN-1 have distinct roles downstream of p38/PMK-1. SKN-1 mediates responses to oxidative stress by regulating classical phase II detoxification genes, whereas ATF-7 governs immune responses, such as resistance to *P. aeruginosa* (Foster et al., 2020; Zhu et al., 2022). Although *mul-1* induction was strongly reduced in *atf-7* mutants, a residual response remained detectable, suggesting that additional factors may contribute to *mul-1* activation in parallel with ATF-7 (Fig. 4E-F). Mutations in the *daf-2* insulin receptor and the *daf-16* transcription factor show no effect on *mul-1* induction (Fig. S4).

**Figure 4.**
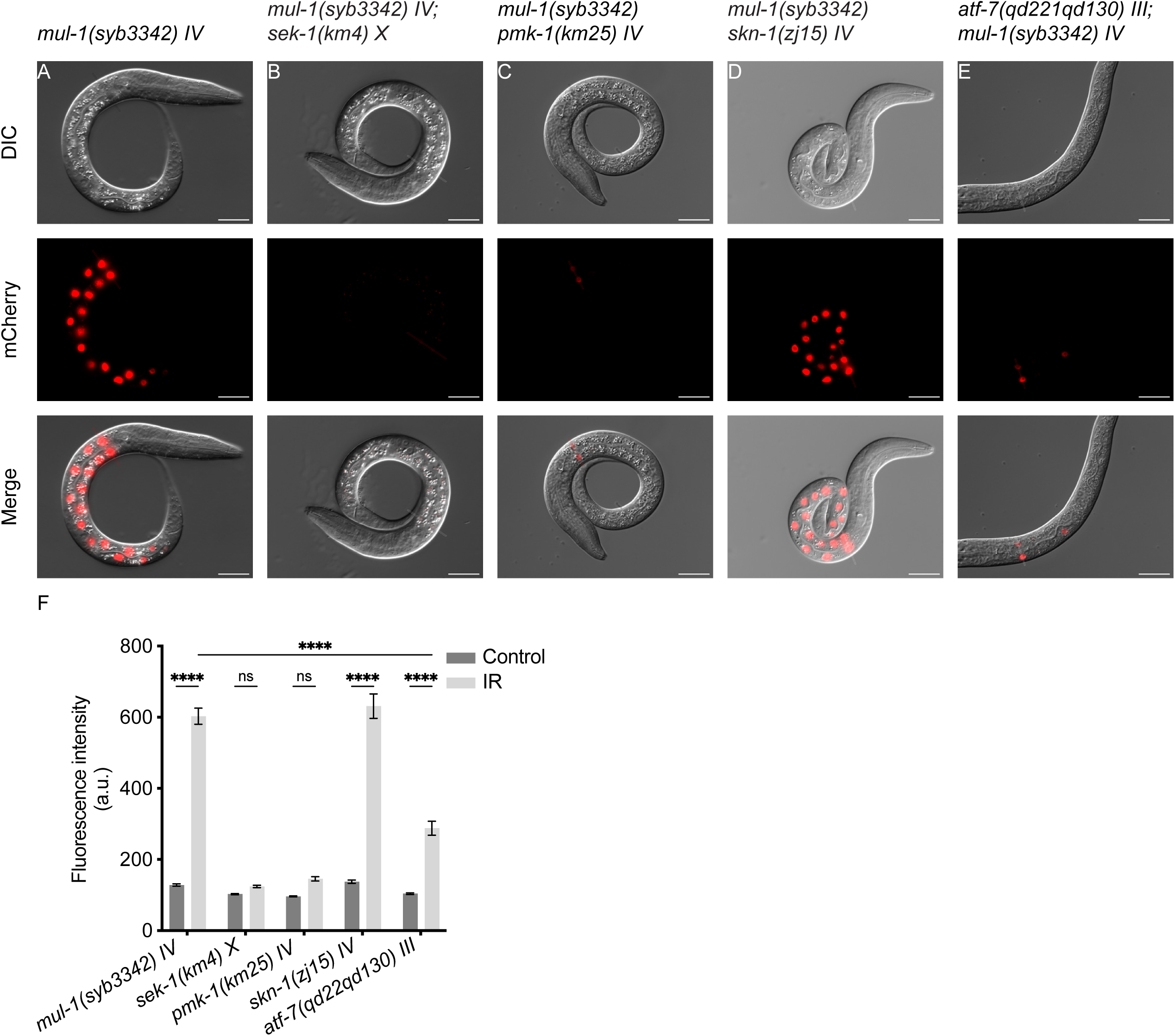
The p38 MAPK pathway and ATF-7, but not SKN-1, are required for mul-1 induction following IR. (A) IR-induced *mul-1* reporter expression in *mul-1(syb3342) IV* animals. (B-C) Loss of the core p38 MAPK components *sek-1* and *pmk-1* abolishes *mul-1* induction in response to IR, indicating an essential role for this signaling pathway. (D-E) Genetic analysis of downstream transcription factors shows that *atf-7*, but not *skn-1*, is required for IR-dependent *mul-1* activation. (F) Quantification of fluorescence intensity in anterior intestinal nuclei across genotypes and conditions. Animals were treated at the L1 stage. Scale bars, 20 µm. Data are shown as mean ± SEM and include animals from at least three independent experiments (n = 20-30 animals per condition). Statistical analysis was performed using two-way ANOVA followed by Tukey’s multiple comparisons test.

To visualize MUL-1 protein, we created a translational reporter, *mul-1::linker::eGFP(gt3545)*. Detecting MUL-1 was challenging due to autofluorescence from gut granules, which interfered with signal clarity. To overcome this, we introduced the *glo-1(zu391)* mutation, which disrupts gut granule formation, thus reducing autofluorescence without affecting intestinal function (Hermann et al., 2005) (Fig. 5A). Under control conditions, MUL-1 expression was not detectable in the *mul-1(gt3545); glo-1(zu391)* reporter strain (Fig. 5B, F, G-H). After IR and H_2_O_2_ treatment, MUL-1 expression was induced, resulting in a low but visible diffuse cytoplasmic signal in intestinal cells, along with distinct cytoplasmic puncta (Fig. 5C, D, F), MUL-1 expression was increased in *prdx-2(gk169); mul-1(gt3545); glo-1(zu391)* animals, indicating that endogenous oxidative stress promotes MUL-1 induction (Fig. 5I-J) in line with the transcriptional induction (Fig. 3I, J, L). An intense cytoplasmic signal can be observed in a high copy MUL-1::eGFP transgene (Fig. 5E) (Hoffman et al., 2020).

**Figure 5.**
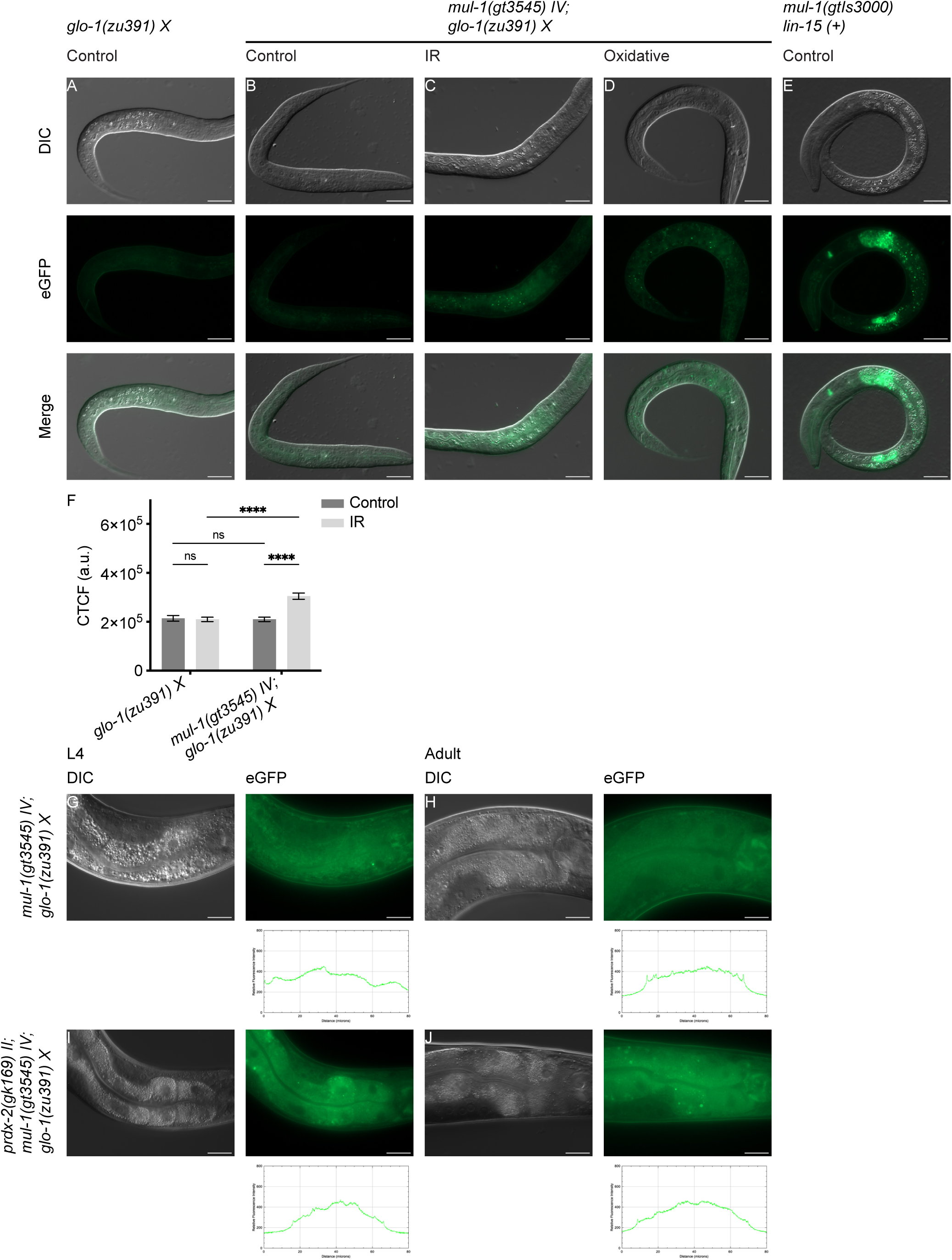
MUL-1 protein accumulates in response to IR and oxidative stress. (A) The *glo-1(zu391) X* mutation reduces intestinal autofluorescence, improving visualization of fluorescent reporters. (B) No detectable MUL-1 expression is observed in *mul-1(gt3545) IV; glo-1(zu391) X* animals under control conditions. (C-D) Following exposure to IR or H_2_O_2_ treatment, MUL-1 protein accumulates in intestinal cells, displaying diffuse cytoplasmic localization and formation of cytoplasmic puncta. (E) An integrated multicopy *mul-1::eGFP* transgene displays detectable basal expression in intestinal cells under control conditions, with cytoplasmic and punctate localization, particularly evident in anterior intestinal cells. (F) Quantification of intestinal MUL-1 fluorescence intensity following IR exposure, measured as corrected total cell fluorescence (CTCF). (G-H) Basal MUL-1 protein levels in *mul-1(gt3545) IV; glo-1(zu391) X* animals at the L4 and adult stages, intestinal MUL-1 protein signal is not readily detectable under control conditions. (I-J) In contrast, *prdx-2(gk169) II; mul-1(gt3545); glo-1(zu391) X* animals display detectable intestinal MUL-1 protein accumulation at the L4 and adult stages, with cytoplasmic distribution and punctate structures. Unless otherwise indicated, animals were analysed at the L1 stage. Scale bars, 20 µm. Data are shown as mean ± SEM from at least three independent experiments (n = 20-30 animals per genotype and condition). Statistical analysis was performed using two-way ANOVA followed by Tukey’s multiple comparisons test.

### MUL-1 mitigates oxidative stress and modulates DNA damage-induced germ cell apoptosis

We could not identify an overt phenotype associated with *the mul-1* reporter line lacking the open reading frame under basal conditions, based on progeny numbers, embryonic lethality, or lifespan (Fig. 6A-C). Additionally, after exposure to IR, we did not observe any deviation from wild type in developmental progression from the L1 stage or in progeny survival at the L4 stage (see below, Fig. 7G). To further explore the role of *mul-1* in oxidative stress management, we used the CellROX Green assay. This fluorescent probe detects multiple ROS, including H_2_O_2_, superoxide, and hydroxyl radicals (Palacin-Martinez et al., 2024). Under control conditions, both wild-type and *mul-1(syb3342)* mutants showed minimal green fluorescence, indicating low basal ROS levels (Fig. 6D, E, G). Following IR exposure, wild-type animals showed a moderate increase in ROS levels (Fig. 6D, F), whereas *mul-1(syb3342)* mutants exhibited a stronger signal, especially in intestinal cells (Fig. 6D, H). We then directly tested sensitivity to H_2_O_2_ exposure and first identified the most suitable concentration, finding that L1 worms treated with 1 mM H_2_O_2_ developed normally, while those treated with 5 mM H_2_O_2_ were uniformly arrested at the L1 stage; treatment with 2.5 mM resulted in an intermediate response (Fig. S5A). When assessing sensitivity to 2.5 mM H_2_O_2_, we observed that developmental progression was moderately delayed in *mul-1(syb3342)*, comparable to *daf-16* and *pmk-1* mutants, which served as positive controls (Fig. 6I).

**Figure 6.**
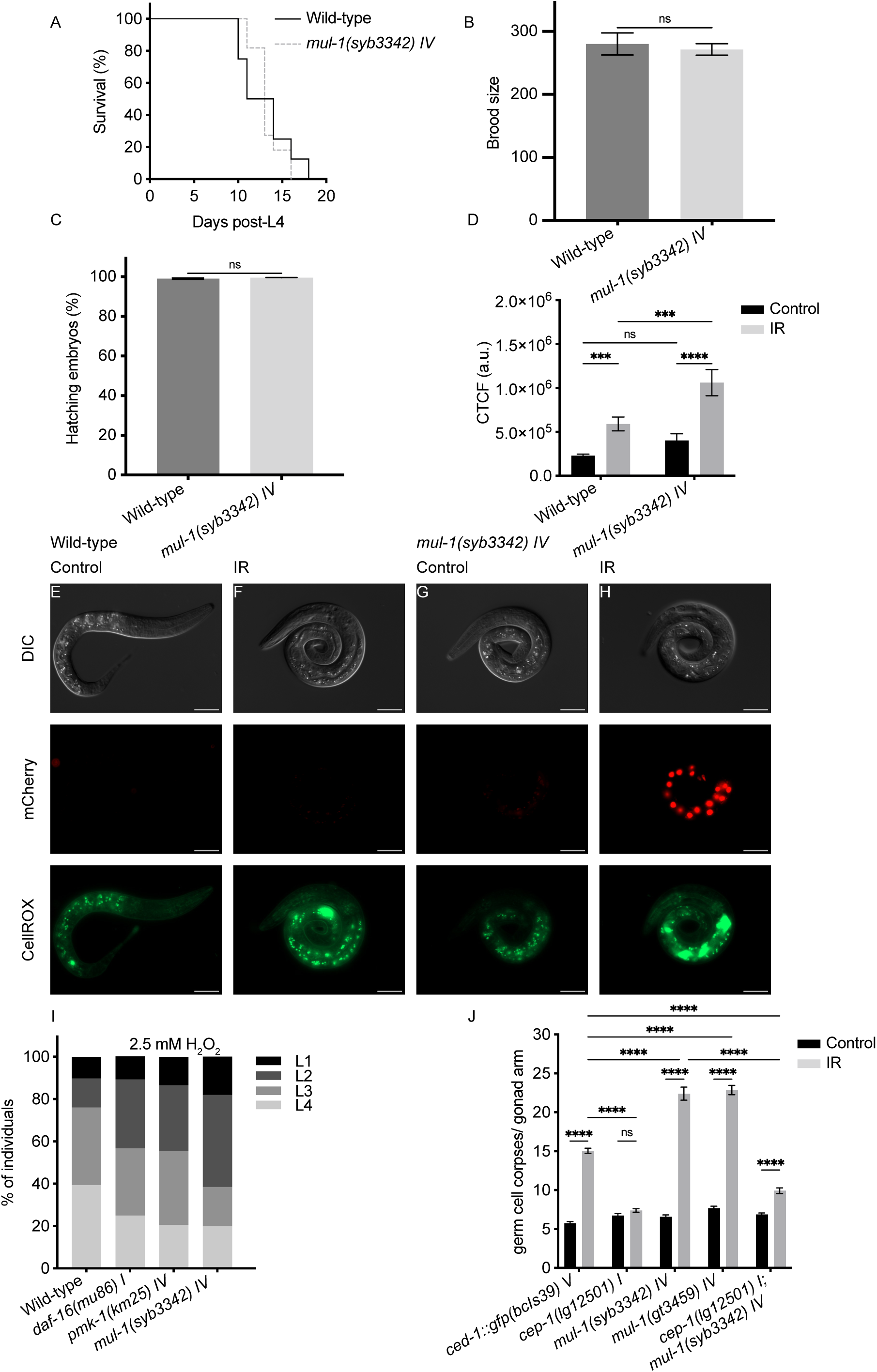
MUL-1 buffers oxidative stress, promotes developmental progression under oxidative stress, and restrains IR-induced germline apoptosis. (A) Lifespan analysis under control conditions reveals no significant difference between wild-type and *mul-1* mutants (n = 10 animals per genotype). (B) Brood size analysis of wild-type and *mul-1(syb3342) IV* animals under control conditions shows no significant difference in total progeny (n = 10-20 animals per genotype). (C) Embryonic viability, assessed by the fraction of hatched embryos, is comparable between wild-type and *mul-1*mutants (n = 10-20 animals per genotype). (D) Quantification of CellROX Green fluorescence intensity in wild-type and *mul-1(syb3342) IV* animals under control conditions and following IR reveals exaggerated ROS accumulation in *mul-1* mutants after IR (n = 20-30 animals per genotype and condition). (E-F) Representative CellROX green images of wild-type animals show low basal ROS levels under control conditions and a moderate increase following IR. (G-H) In contrast, *mul-1(syb3342) IV* animals display low basal CellROX signal under control conditions, but accumulate excessive ROS after IR, with strong fluorescence particularly evident in intestinal cells. (I) Developmental stages distribution 48 h after exposure to 2.5 mM H_2_O_2_ at the L1 stage (n = 60 animals per genotype). Developmental stages were scored based on vulval morphology and overall body size. *mul-1 (syb3342) IV* mutants display a moderate delay in developmental profession compared to wild-type animals, comparable to that observed in *daf-16* and *pmk-1* mutants, which were included as positive controls for oxidative stress sensitivity. (J) Quantification of germ cell corpses 24 h after exposure to IR reveals a hyperinduction of apoptosis in *mul-1* mutants, which is suppressed in *cep-1; mul-1* double mutants (n = 20-30 animals per genotype and condition). This phenotype was independently confirmed using a *mul-1(STOP-IN)* null allele. Germ cell corpses were scored using the *ced-1::gfp*reporter. L1 animals were analysed unless otherwise indicated. Scale bars, 20 µm. Data are shown as mean ± SEM from at least three independent experiments. Lifespan (A) was analyzed using log-rank (Mantel-Cox) and Gehan-Breslow-Wilcoxon tests. Brood size and embryonic viability (B-C) were analyzed using unpaired two-tailed Student’s *t*-tests. CellROX fluorescence (D) and germ cell apoptosis (J) were analyzed by two-way ANOVA followed by Tukey’s or Šidak’s multiple comparisons tests.

**Figure 7.**
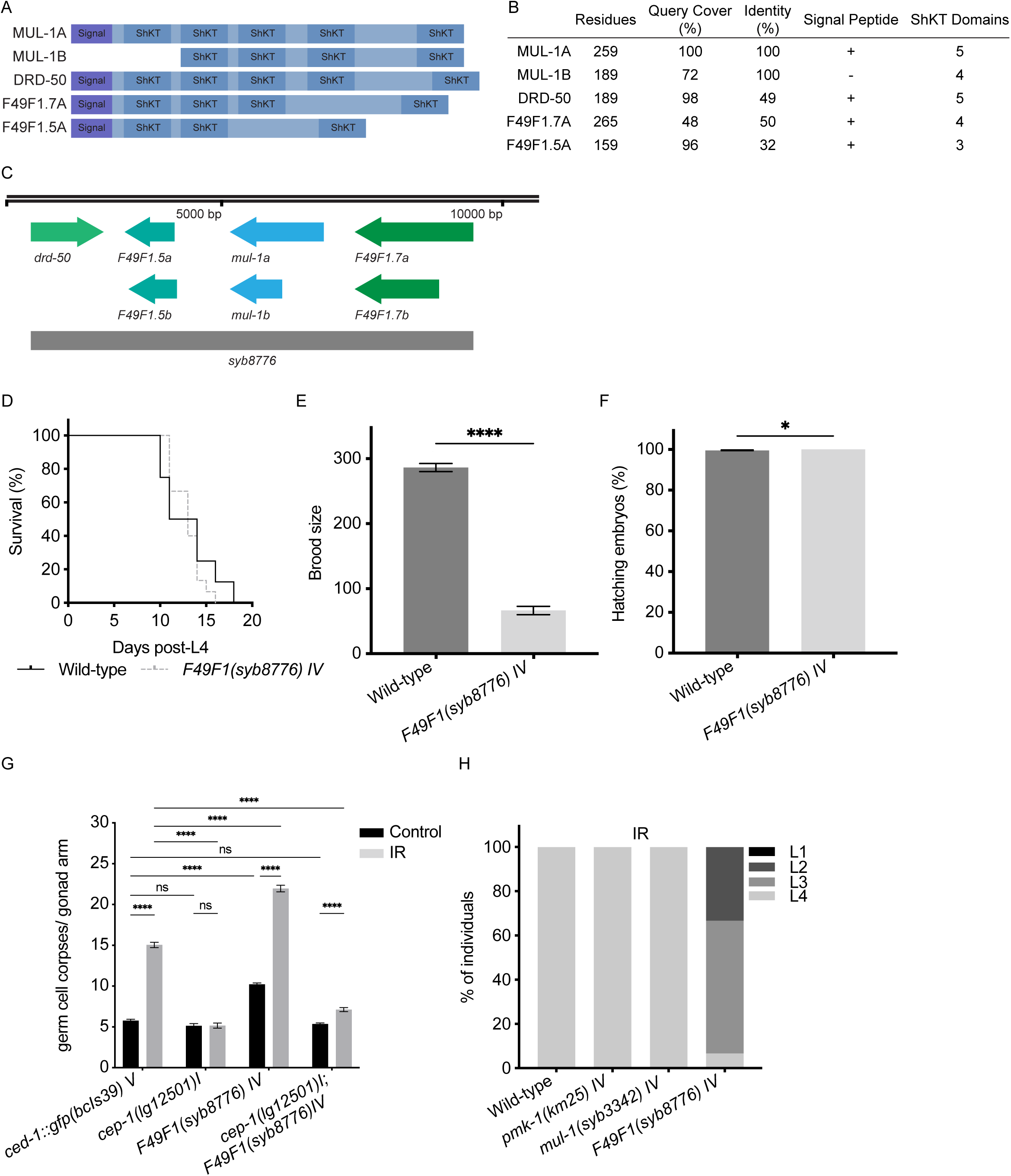
A ShKT-containing MUL-1 paralog cluster acts redundantly to maintain germline homeostasis and enable developmental recovery after IR. (A) Schematic representation of the domain architecture of MUL-1 and its closest paralogs, all encoding ShKT domain-containing proteins. (B) Pairwise sequence identity matrix comparing MUL-1 and related paralogs. (C) Schematic representation of the *F49F1* genomic locus on chromosome IV showing the organization of *mul-1* and its paralogs. The *syb8776* allele corresponds to a deletion of an approximately 7.9 kb region encompassing all four genes. (D) Lifespan analysis of *F49F1* quadruple mutant shows no significant difference compared to wild-type under control conditions (n = 10-20 animals per genotype). (E) Progeny production is significantly reduced in the *F49F1* quadruple mutant (n = 10-20 animals per genotype). (F) Embryonic lethality remains unchanged in *F49F1* quadruple mutants (n = 10-20 animals per genotype). (G) Germ cell apoptosis is elevated in the *F49F1* quadruple mutants under both control and IR conditions (n = 20-30 animals per condition). (H) Developmental stage distribution of animals exposed to IR at the L1 stage and scored after 48 hours recovery (n = 60 animals per genotype). Data are shown as mean ± SEM. Lifespan analyses (D) were performed using log-rank (Mantel-Cox) and Gehan-Breslow-Wilcoxon tests. Progeny production and embryonic viability (E-F) were analyzed using unpaired two-tailed Student’s *t*-tests. Germ cell apoptosis (G) was analyzed by two-way ANOVA followed by Tukey’s multiple comparisons test.

Next, we analyzed germ cell apoptosis using a widely used reporter where the CED-1 apoptotic corpse receptor is tagged with GFP (Zhou et al., 2001). Under basal conditions, *mul-1(syb3342)* animals exhibited normal levels of apoptosis (Fig. 6J, lanes 1 and 5). In contrast, following IR, apoptosis was hyperinduced in *mul-1(syb3342),* a finding confirmed when using a second *mul-1* allele (Fig. 6J, lanes 2, 6, 8). Excessive IR-dependent apoptosis was suppressed in *cep-1(lg12501)*, which is defective for the nematode p53-like transcription factor (Fig. 6J, lanes 4 and 10). Complementation analyses using *mul-1::eGFP* and *mul-1::3xHA* alleles under the same conditions confirmed that both tagged MUL-1 proteins are functional, as they suppress the extra apoptosis phenotype of *mul-1* to wild-type apoptosis levels (Fig. S5B).

### MUL-1 belongs to a conserved ShKT-containing gene cluster that modulates oxidative stress and germline homeostasis

We aimed to investigate if *mul-1* might act redundantly. Performing BLAST searches for MUL-1 paralogs and scanning through the WormBase we indeed found multiple MUL-1 paralogs. *mul-1* is part of a four-gene cluster on chromosome IV that includes three additional closely related genes (*drd-50*, F49F1.5, F49F1.7) (Fig. 7A, B, Suppl. Table 1). MUL-1 paralogs contain multiple ShKT domains with six conserved cysteines, forming three disulfide bonds that stabilize a compact double α-helix structure (Fig. 7A). This clustering suggests potential co-regulation or shared functions. Consistent with this, transcriptional analysis revealed a strong induction of *mul-1* upon IR, whereas its paralogs display only modest changes, indicating differential regulation within the cluster (Fig. S6A). We refer to the *syb8776* mutation when taking out all 4 paralogs as the ‘quadruple mutant’ (Fig.7C). To test for redundancy, we started by analyzing the quadruple mutant and focused on developmental progression and lifespan, a key measure of organismal resilience affected by genes involved in stress response, genomic stability, and cellular homeostasis.

While wild-type or single deletions of *mul-1* or *pmk-1* do not impair developmental progression following IR, the quadruple mutant exhibits delayed larval development, indicating functional redundancy among MUL-1 paralogs during recovery from genotoxic stress.maintenance (López-Otín et al., 2023). The quadruple mutant showed no change in lifespan compared to wild-type (Fig. 7D). However, the quadruple mutant had a significant reduction in progeny (Fig. 7 E). In contrast, no embryonic lethality was observed in the quadruple mutant, similar to the wild-type (Fig. 7F). Importantly, expression of the multicopy *mul-1(gtIs3000)* transgene restored brood size in the quadruple mutant without affecting embryonic survival (Fig. S6B-C). We then tested if reduced progeny in the quadruple mutant was correlated with excessive germ cell apoptosis and found that it was the case, both with and without IR treatment (Fig. 7G). Excessive apoptosis was CEP-1/p53 dependent under both conditions. Given the redundancy of MUL-1-like proteins, we tested whether larval development is delayed upon treating L1 stage animals with IR, and found that this is true for the quadruple mutant (Fig. 7H). In contrast, single deletions of *mul-1* or *pmk-1* do not impair developmental recovery after IR, the combined loss of *mul-1* paralogs slows development.

### MUL-1 paralogs protect from *Pseudomonas aeruginosa* infection

We hypothesized that MUL-1 and its closely related paralogs may protect against bacterial infection, commonly associated with oxidative stress. Therefore, we tested susceptibility to *P. aeruginosa* PA14 infection, a model commonly used in vertebrates and *C. elegans* (Tan et al., 1999), and known to induce oxidative stress in the nematode (Zhang et al., 2025). We observed that *mul-1(syb3242)* animals died slightly earlier than wild type, with the quadruple mutant being the most sensitive, comparable to the *pmk-1* positive control (Fig. 8A). *mul-1* expression was robustly induced upon exposure to PA14, as indicated by increased reporter fluorescence intensity relative to OP50-fed controls (Fig. 8B-D).

**Figure 8.**
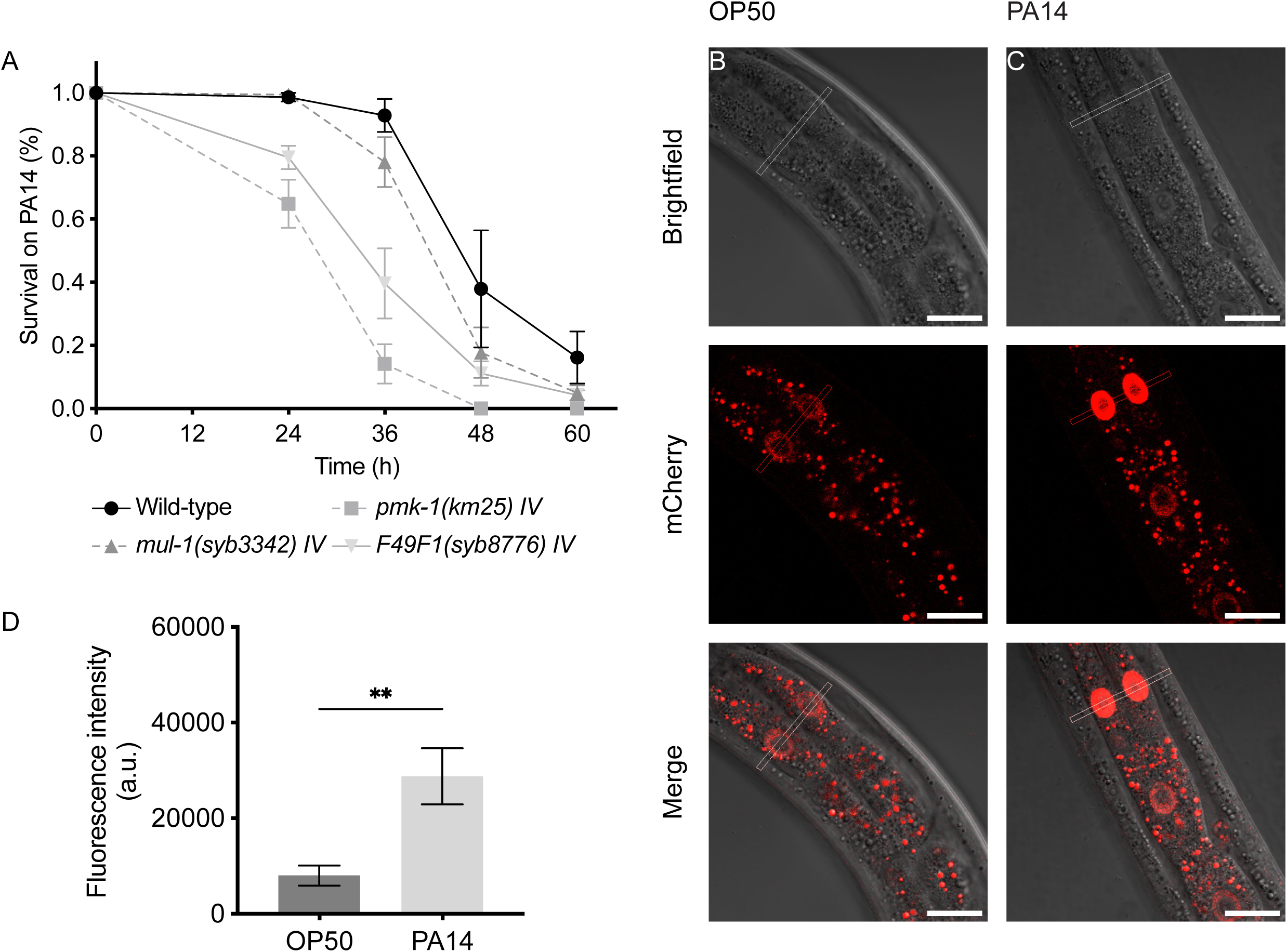
Redundant MUL-1 paralogs contribute to resistance against *P. aeruginosa* infection. (A) Survival analysis of wild-type animals, *mul-1(syb3342) IV*, *pmk-1* and the *F49F1* mutants following exposure to *P. aeruginosa* PA14. While *mul-1* single mutants do not display increased sensitivity, the quadruple mutant exhibits reduced survival comparable to the *pmk-1* positive control (n = 60 animals per genotype). (B-C) Representative images showing reporter expression in adult animals exposed to *E. coli* OP50 (B) or *P. aeruginosa*PA14 (C). (D) Quantification of fluorescence intensity reveals robust induction of *mul-1* reporter expression upon PA14 exposure compared to OP50-fed controls. Data are shown as mean ± SEM from three independent experiments. Fluorescence intensity (D) was analyzed using an unpaired two-tailed Student’s *t*-test with Welch’s correction.

### Loss of MUL-1 paralogs activates oxidative stress response pathways via *daf-16*/FoxO and *skn-1*/Nrf2

If MUL-1-like proteins act by scavenging or sensing oxidative stress, their absence might lead to increased endogenous ROS and possibly activate compensatory stress-response pathways. We thus tested if ROS is induced in the quadruple mutants even in the absence of IR, and found that this is the case using the CellROX green assay (Fig. 9A-C).

**Figure 9.**
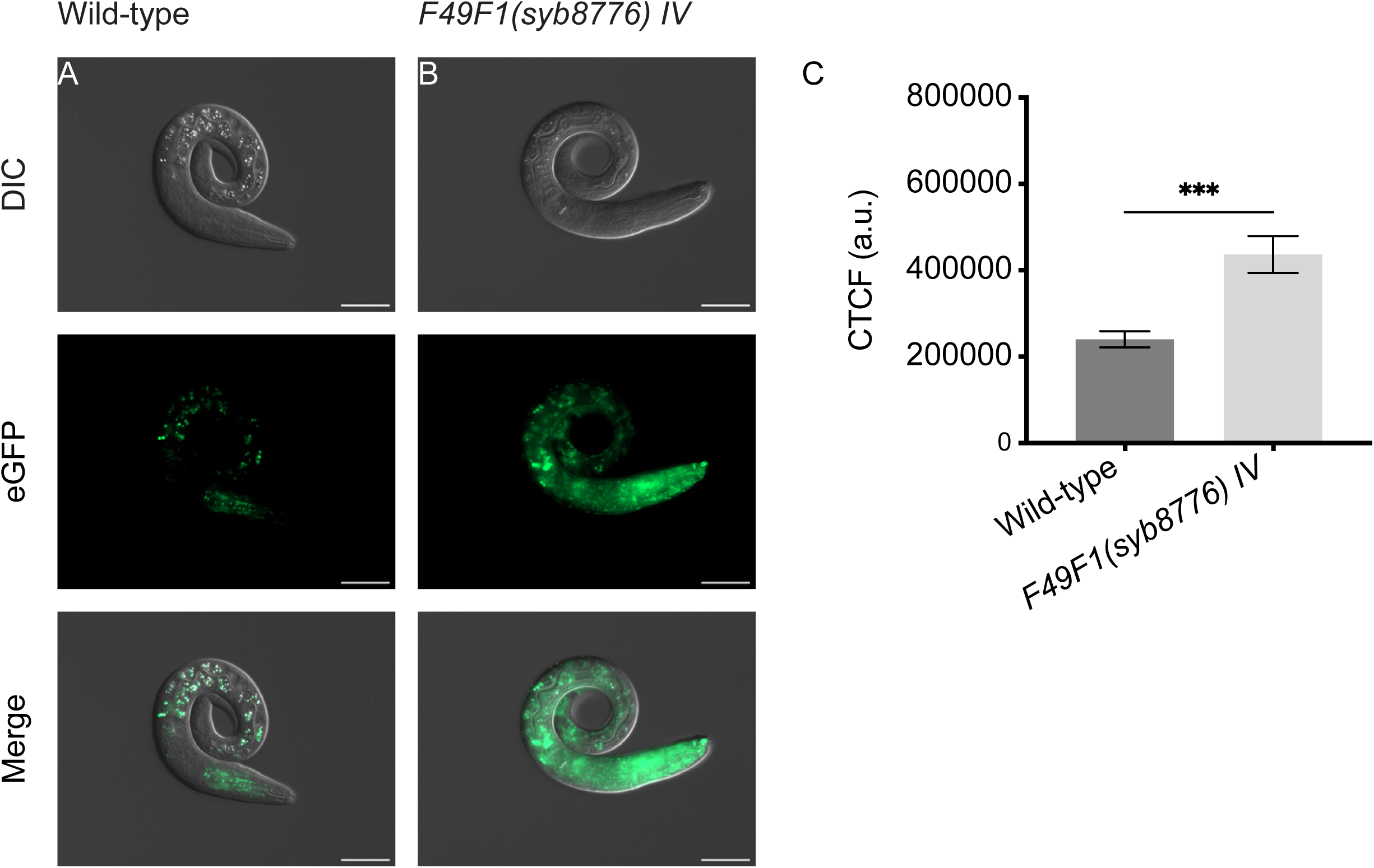
MUL-1 paralogs restrain basal ROS accumulation. Representative images of wild-type (A) and *F49F1(syb8776) IV* animals (B) stained with CellROX Green under control conditions. (C) Quantification of fluorescence intensity reveals increased basal ROS levels in the quadruple mutant compared to wild-type animals (n = 20 animals per genotype). Data are shown as mean ± SEM from at least three independent experiments. Statistical analysis was performed using an unpaired two-tailed Student’s *t*-test with Welch’s correction.

To examine if this leads to the activation of compensatory pathways, we investigated oxidative stress response pathways regulated by *daf-16*/FoxO and *skn-1/Nrf*2 (Doonan et al., 2008; Inoue et al., 2005; Leiers et al., 2003). We generated translational reporters for *sod-3* and *gst-4* to assess pathway activation by fusing eGFP to *sod-*3 *(sod-3(gt3598))* and mCherry to *gst-*4 *(gst-4(gt3596))*. We combined both reporters with the *glo-1(zu391)* mutation to reduce autofluorescence from intestinal gut granules (Hermann et al., 2005). *sod-3* encodes a mitochondrial manganese superoxide dismutase (MnSOD), which neutralizes superoxide radicals and is often linked to increased stress resistance and longevity (Doonan et al., 2008). GST-4 is a phase II detoxification enzyme, glutathione S-transferase, which has a peroxidase activity and catalyzes the conjugation of glutathione (GSH) to electrophilic compounds, promoting detoxification and excretion (Hurst et al., 1998; Inoue et al., 2005; Leiers et al., 2003). Analysis of GST-4::mCherry and SOD-3::eGFP expression in wild-type L1 larvae confirmed that GST-4 is primarily expressed in the intestine, with additional localization in head hypodermal cells (Fig. 10A). In contrast, SOD-3::eGFP fluorescence was localized to the pharynx, especially in the anterior bulb, with a faint but detectable signal around the terminal bulb of the pharynx. No *sod-3* expression was observed in the intestine under normal conditions, nor in the hypodermis, body wall muscles, neurons, or tail (Fig. 10A). Deletion of *mul-1* alone did not significantly change GST-4 expression but increased SOD-3 expression in the pharynx (Fig. 10B). Remarkably, the *F49F1(syb8776)* quadruple deletion caused a strong induction of GST-4 throughout the body, especially in the anterior gut, with widespread upregulation of SOD-3 (Fig. 10C). Consistent with these findings, *prdx-2* mutants, which are known to accumulate endogenous ROS and which we show to induce *mul-1* (Fig. 3 and 5), also showed strong GST-4 induction and pharyngeal SOD-3 expression (Fig. 10D). As expected, GST-4 induction in *prdx-2* mutants was *skn-1*-dependent (Fig. 10E), while SOD-3 induction required *daf-16* (Fig. 10F). Overall, these results suggest that deleting *mul-1* and its paralogs increases oxidative stress and the expression of key genes involved in oxidative stress response.

**Figure 10.**
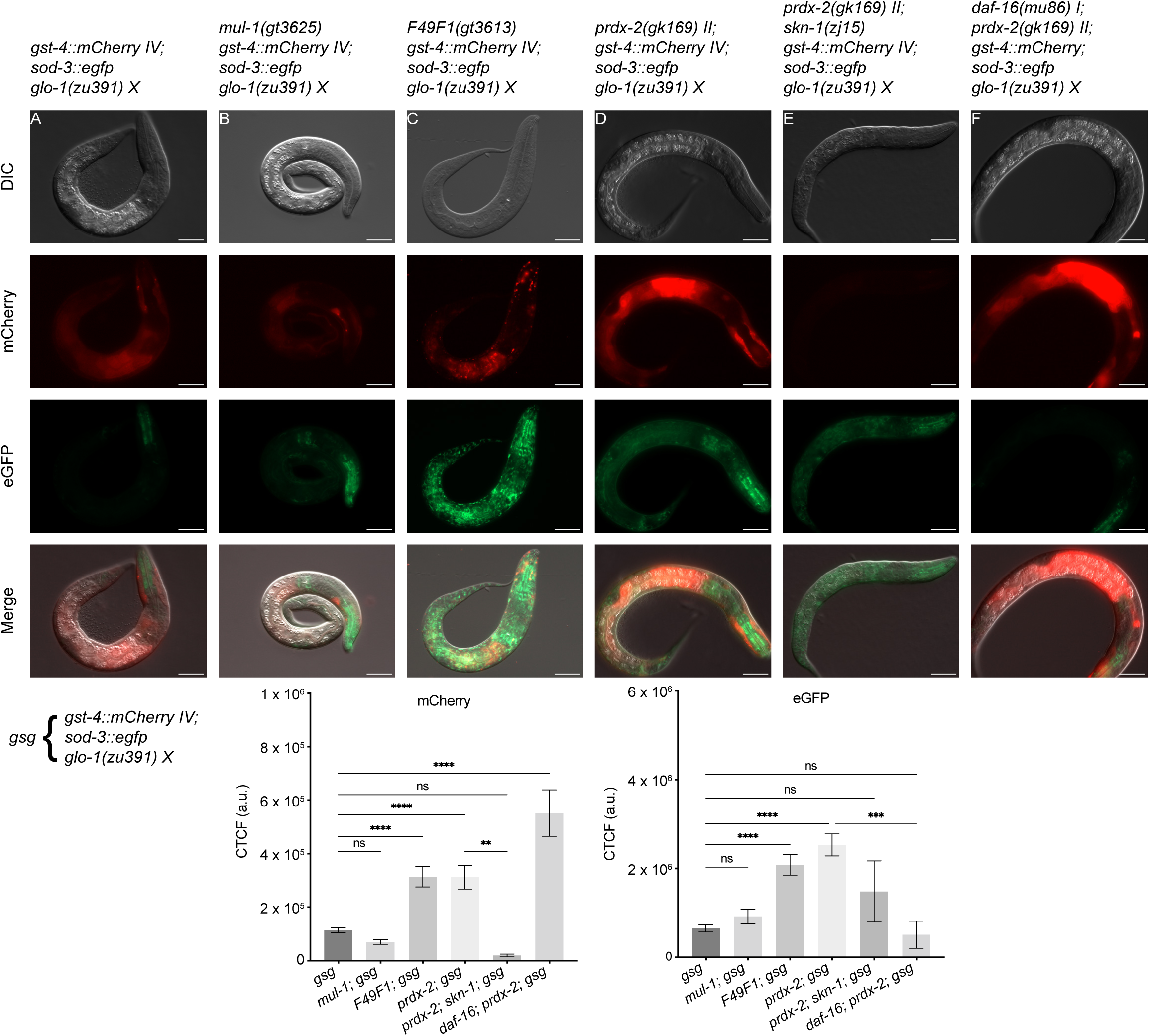
Compensatory induction of GST-4::mCherry and SOD-3::eGFP. (A) Expression pattern of the oxidative stress reporters *gst-4::mCherry* and *sod-3::eGFP* in wild-type L1 larvae carrying the *glo-1(zu391) X* mutation. *gst-4::mCherry* is predominantly expressed in the intestine, whereas *sod-3::eGFP* localizes mainly to the pharynx. (B) Deletion of *mul-1* alone does not alter *gst-4* or *sod-3* expression. (C) The quadruple mutant shows strong induction of *gst-4* throughout the body, particularly in the anterior intestine, together with widespread upregulation of *sod-3*. (D) *prdx-2(gk169) II* mutants exhibit robust induction of *gst-4* and increased *sod-3* expression, consistent with elevated endogenous oxidative stress. (E) Induction of *gst-4* in *prdx-2* mutants requires *skn-1*. (F) Induction of *sod-3* in *prdx-2* mutants is abolished in the absence of *daf-16*. Representative images are shown. Quantification of *gst-4::mCherry* and *sod-3::eGFP* fluorescence was performed on 20-30 animals per genotype per experiment. Data are shown as mean ± SEM from at least three independent experiments. Statistical analysis was performed using one-way ANOVA followed by Tukey’s multiple comparisons test.

### Genetic interaction with further MUL-1 paralogs

Our results are consistent with MUL-1 proteins acting redundantly to protect animals from oxidative stress. Given that the nematode genome encodes multiple additional MUL-1 paralogs we wanted to test this notion more generally and examined a more distantly related MUL-1 paralog SYSM-1, given its reported role as an apoptosis effector, deficiency leading to decreased, and not increased DNA damage induced germ cell apoptosis. SYSM-1 acts cell-non autonomously, being secreted from the gut and functioning independently of p53 (Soltanmohammadi et al., 2022). We confirmed that *sysm-1* mutants are defective for DNA damage-induced apoptosis (Soltanmohammadi et al., 2022) (Fig. 11A). However, analysis of *sysm-1; mul-1* double mutants as well as quadruple mutant in conjunction with *sysm-*1 revealed that the excessive apoptosis phenotype observed in *mul-1* single mutants as well as in the quadruple mutant where excessive apoptosis occurs even without IR is not suppressed by *sysm-1.* In other words, the apoptosis defect associated with *sysm-1* is bypassed by *mul-1* and its paralogs (Fig. 11A).

**Figure 11.**
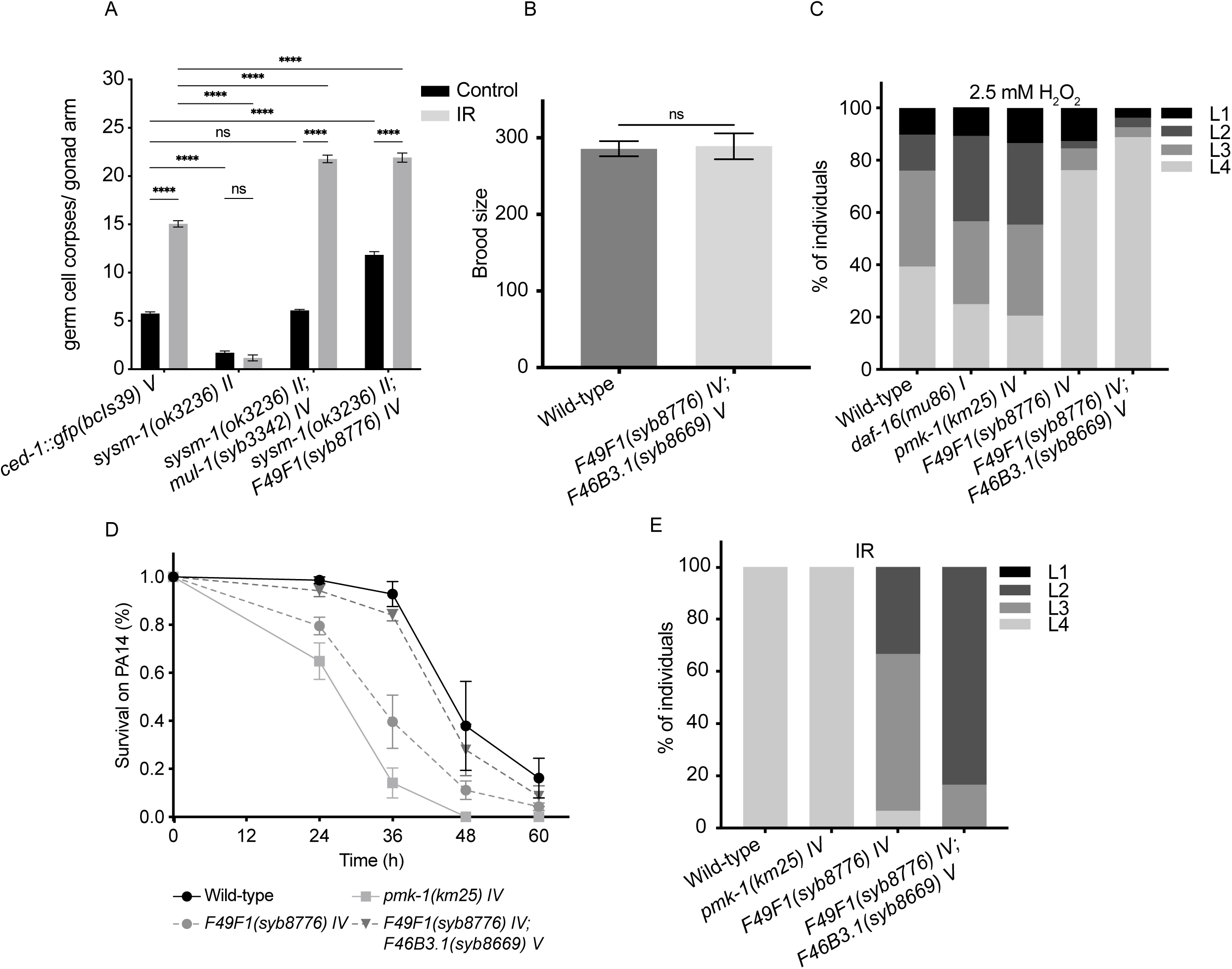
Genetic interactions between MUL-1 and its paralogs shape organismal responses to oxidative and genotoxic stress. (A) Suppression of apoptosis defect of *sysm-1*. (B) Genetic interaction with the F46B3.1 MUL-1 paralog. (C) Suppression of H_2_O_2_ sensitivity by F46B3.1 loss-of-function. (D) Suppression of *Pseudomonas* PA14 sensitivity by F46B3.1 loss-of-function. (E) *F46B3.1* loss-of-function increases sensitivity to IR. Data are shown as mean ± SEM from at least three independent experiments. Germ cell apoptosis (A) was analyzed by two-way ANOVA followed by Tukey’s multiple comparisons test (n = 20-30 animals per condition). H_2_O_2_ and IR sensitivity assays (C, E) were analyzed by two-way ANOVA (n = 60 animals per genotype). Survival assays (D) were analyzed using log-rank (Mantel-Cox) tests (n = 60 animals per genotype).

We next generated and analysed *syb8669* a deletion of F46B3.1, the most closely related MUL-1 paralog located outside the MUL-1 paralog cluster (quadruple mutant). Double mutant analysis of the allele leads to complex genetic interactions with *mul-1* and the quadruple mutant: While the single mutant has no overt phenotype compared to N2, the quintuple mutant suppressed some phenotypes, while enhancing others. *syb8669* suppressed the reduced fecundity of the quadruple mutant (Fig.11B). Conversely, the reduced H_2_O_2_ sensitivity of the quadruple mutant was further suppressed to an extent such that quintuple mutants are partially resistant compared to WT (Fig. 11C). Also, the hypersensitivity towards *Pseudomonas* PA14 of the quadruple mutant is suppressed in the quintuple mutant (Fig. 11D). In contrast, quintuple mutants are hypersensitive to IR (Fig. 11E). All in all, the experiments including *sysms-1* and *syb8669* point towards a more complex picture where MUL-1 paralogs can have opposing functions, in line with the hypothesis that MUL-1 like proteins might besides being scavengers may also act as rheostates for ROS.

## Discussion

We initiated our study by focusing on MUL-1 and later included the four closest paralogs in our analyses. MUL-1 and its close paralogs are unstructured, except for possessing 3-5 ShKT domains. These domains are cysteine-rich motifs initially described in sea anemone toxins and are widely found in invertebrate proteins, although their function in nematodes remains largely unexplored (Rangaraju et al., 2010; Sachkova et al., 2020). We postulate that nematode multi-ShKT domain proteins may act as scavengers or rheostats of oxidative stress, owing to their potential to scavenge ROS via disulfide formation facilitated by the six cysteines in each ShKT domain (Fig. 12, see below).

**Figure 12.**
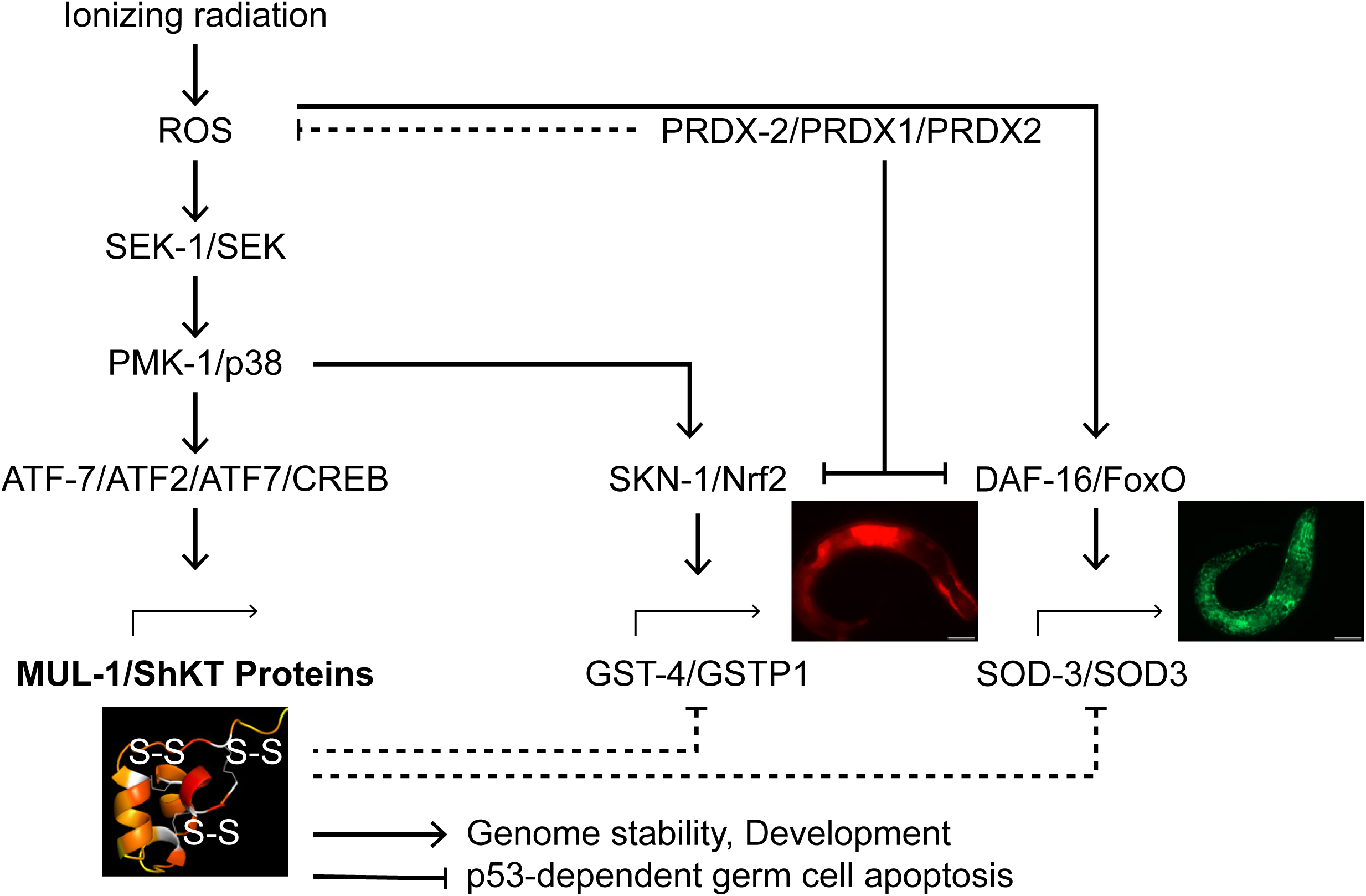
Model for the integration of DNA damage and redox signaling by MUL-1 and ShKT domain proteins. IR increases ROS, engaging the SEK-1/PMK-1 p38 MAPK pathway and its downstream transcription factor ATF-7 to induce MUL-1 and other ShKT-domain proteins. Under physiological conditions, PRDX-2 limits basal ROS levels. Upon stress, ShKT proteins function as redox-responsive modulators that limit the magnitude of antioxidant gene activation. In parallel, elevated ROS activate SKN-1/Nrf2 to promote GST-4 expression and DAF-16/FoxO to drive SOD-3 expression. In the absence of ShKT proteins, derepression of GST-4 and SOD-3 enhances antioxidant capacity, thereby supporting genome stability and normal development while mitigating p53/CEP-1-dependent germ cell apoptosis.

We certainly do not rule out that MUL-1 and its paralogs are mucin-like proteins (Hoffman et al., 2020). MUL-1 encodes a 42-amino-acid domain highly enriched in serine/threonine residues, which is akin to mammalian mucins that are highly enriched for serine/threonine throughout most of their length. The parasitic nematode *Toxocara canis* encodes four secreted proteins, each with an N-terminal signal peptide for secretion and an 83-97 amino acid S/T-enriched mucin domain, N-terminal to two ShkT domains (Loukas et al., 2000). For MUL-1, prominent enrichment of serine and threonine residues, which occurs along the entire length of mammalian mucins comprising several thousand amino acids (Johansson et al., 2013), occurs only in a 44-amino-acid unstructured region between the 4th and 5th ShKT domains. Irrespective, our combined genetic analysis indicates that MUL-1 and related paralogs are involved in a circulatory regulatory circuit associated with oxidative stress as discussed below.

We discovered that the transcriptional induction of MUL-1 by IR, which generates ROS, is restricted to the gut, with the most pronounced effects observed in the anterior cells. Notably, MUL-1 induction is not triggered by DNA-damaging agents such as the methylating agent MMS or the DNA cross-linking agent cisplatin, nor by osmotic stress or starvation, but rather by oxidative stress, as demonstrated by direct exposure to H_2_O_2_ or increased endogenous H_2_O_2_ levels in peroxidase-deficient *prdx-2* mutants*. mul-1* induction is mediated by the p38 MAPK signaling pathway, consistent with previous RNAi-based studies (Kimura et al., 2012), and requires the transcription factor ATF-7, but not SKN-1. While SKN-1 is widely recognized as a master regulator of oxidative stress responses by inducing phase II detoxification genes (Blackwell et al., 2015; Foster et al., 2020), our findings underscore a previously underappreciated role for ATF-7 in orchestrating transcriptional responses to ROS accumulation. At first glance, gut expression seems intriguing; however, it aligns with bacterial-nematode infections, in which bacterial pathogens and *C. elegans* produce ROS upon pathogen exposure *(Chavez et al., 2007; Hoeven et al., 2011; Jansen et al., 2002; Miranda-Vizuete & Veal, 2017)*. Indeed, *Rhizobium* infection and the associated oxidative stress led to defective genome integrity during larval gut development, resulting in excessive DNA bridges and karyokinesis defects in gut nuclei, with the phenotype most prominent in the anteriormost gut cells (Kniazeva & Ruvkun, 2019). We did not find decreased susceptibility to *P. aeruginosa* infection in the *mul-1* single mutant, as previously reported (Hoffman et al., 2020), but did find increased sensitivity in the quadruple mutant. This is due to us using 5-fluoro-2’-deoxyuridine (FUDR) to prevent germ cell proliferation, animals otherwise producing embryos that hatch inside their parents (bagging phenotype) leading to lethality (Kwon et al., 2024).

Functionally, in our study, *mul-1* mutants exhibited a small increase of ROS after IR, a modest delayed development under oxidative stress conditions, and elevated CEP-1-dependent germ cell apoptosis. *mul-1* is part of a cluster of three additional paralogs, each encoding three to five ShKT domains. Deleting these three paralogs, along with *mul-1* (quadruple mutant), reduced progeny numbers and exhibited CEP-1/p53-dependent germline apoptosis even without irradiation. Also, genetic interactions with SYSM-1, a small, 99-amino-acid, unstructured protein that carries two ShKT domains and is induced by IR (Soltanmohammadi et al., 2022), yielded surprising results. Like MUL-1, SYSM-1 is a transcriptional target of p38 signaling and the downstream ATF-7 transcription factor (Soltanmohammadi et al., 2022). In contrast to MUL-1, which reduces CEP-1 p53-induced germ cell apoptosis, SYSM-1 is essential for DNA damage-induced apoptosis. We find that radiation-induced apoptosis of *sysm-1* mutants is bypassed by *mul-1* single and quadruple mutants. SYSM-1 was suggested to be secreted from the gut to facilitate DNA damage-induced apoptosis in the germ line (Soltanmohammadi et al., 2022).

We show that in the absence the 4 MUL-1 paralogs, endogenous ROS accumulates, and this aligns with our observation that DAF-16-dependent SOD-3::eGFP and SKN-1-dependent Cherry::GST-4 are induced in the *mul-1* quadruple paralog mutant. Together, these data support a circular model of redox signaling, in which MUL-1 and MUL-1-like ShKT domain proteins may function as scavengers or rheostats of ROS (Fig. 12). This way, the loss of the MUL-1 cluster may lead to increased oxidative stress and the compensatory activation of stress pathways, conferring increased survival under oxidizing conditions, but is insufficient to protect against apoptosis induction, sensitivity to IR and *Pseudomonas* infection. A localized balance between the expression of various MUL-1 paralogs and the differential activation of compensatory pathways might determine the activity of different stress response pathways. At present, we do not know how signals associated with *mul-1* single and compound mutants are transmitted across worm tissues, especially to the germ line, where CEP-1-dependent apoptosis is induced. Signaling, and this hypothesis remains to be tested, might be conferred via direct translocation of closely related MUL-1 paralogs, as shown for SYSM-1 (Soltanmohammadi et al., 2022). Alternatively, intercellular signalling could be directly mediated by H_2_O_2_ diffusion across plasma membranes mediated by aquaporins (Sies & Jones, 2020).

ShKT domains were initially characterized as potent toxins derived from sea anemones that inhibit mammalian potassium channels (Castañeda et al., 1995; Harvey & Vita, n.d.; Shafee et al., 2019; Tudor et al., 1998). We postulate that ShKT domains may be involved in redox reactions. If so, ShKT domains are used in redox regulation, and their cysteines could be oxidised akin to the 3 amino acid GSH (glutathione) peptide to its oxidized dimeric (GSSG) form. Each ShKT domain contains six cysteine residues, potentially allowing for extensive disulfide bond formation and redox reactivity. Although such a system may appear inefficient, especially if reductive recycling does not occur, it could serve as a buffering mechanism during acute oxidative stress. In this context, the expansion of proteins primarily composed of ShKT domains in nematodes and other invertebrates might reflect an evolutionary strategy to cope with transient yet potentially lethal oxidative insults. Akin, peroxiredoxin ShKT domains might act as direct H_2_O_2_ scavengers or enable thiol oxidation by relaying H_2_O_2_-derived oxidation equivalents to other proteins (Stöcker et al., 2018).

The hypothesis that ShKT domains might be linked to redox reactions is supported by invertebrate redox-active proteins, such as peroxidases and tyrosinases that carry ShKT domains (Rangaraju et al., 2010). For instance, *C. elegans* MLT-7 and SKPO-1, 2, and 3, peroxidases have acquired an N-terminal ShKT domain and are closely related to the human peroxidasin PXDN which lacks a ShKT domain (Thein et al., 2009; Tiller & Garsin, 2014). These proteins have been shown to crosslink collagen and regulate endothelial basement membrane structure and protect against *E. faecalis* infection (Thein et al., 2009; Tiller & Garsin, 2014). Peroxidase reactions use H*_2_*O_2_ to catalyze the oxidation of various substrates, and *C. elegans* peroxidases modify cuticle collagen structure and permeability (Edens et al., 2001; Myllyharju & Kivirikko, 2004; Thein et al., 2009). Beyond peroxidases, ShKT domains are also present in several *C. elegans* tyrosinases-like proteins (TYR-1 through TYR-6), which belong to the type-3 copper enzyme family and are annotated to contain tyrosinases copper-binding domains together with an N-terminal ShKT module. Although the specific biochemical activities of TYR proteins in *C. elegans* remain untested and are inferred primarily from homology, mammalian tyrosinases are well-established type-3 copper oxidoreductases that function through catalytic redox cycling (Pretzler & Rompel, 2024).

Overall, our combined results indicate that MUL-1-like proteins may act as buffers or rheostats for oxidative stress. It remains to be directly tested if and when ShKT domains are oxidised and if this involves disulfide bond formation. Certainly, it is possible, and this remains to be tested, that MUL-1 like proteins have a role in connecting neuronal circuits and gut behavior, where H*_2_*O_2_ has a role in signaling (Jia et al., 2024; Jia & Sieburth, 2021; Liu et al., 2024). Irrespective, the expansion of ShKT domains in nematodes and other invertebrates may facilitate rapid evolutionary adaptation to the various challenges posed by oxidative stress. The expansion of MUL-1 paralogs may also have facilitated different MUL-1 paralogs having overlapping and opposing functions.

### Limitations of the study

Overall, our combined results indicate that MUL-1 is part of a regulon induced by oxidative stress via p38 MAP kinase signaling, and that MUL-1 and its paralogs may act as buffers or rheostats for oxidative stress. It remains to be directly tested if and when ShKT domains are oxidised and if this involves disulfide bond formation. MUL-1 paralog SYSM-1 was previously shown to be secreted from the gut and taken up in the germ line. Analysing high copy MUL-1 we do not see any evidence for germ line localization, but acknowledge that this might be due to the limited sensitivity of GFP. We recognize that we have not investigated if MUL-1 and its paralogs act cell non-autonomously, which will be an interesting future question. Also, our analysis largely depends on the analysis of the *mul-1* single mutant and the quadruple mutant where all 4 paralogs of the locus are deleted. It will be interesting to investigate how all single, double and triple mutant combinations behave relating to apoptosis induction, H_2_O_2_ resistance, the accumulation of ROS as well as the activation of compensatory GST-4 and SOD-3 activation. Finally, we acknowledge that we do not provide direct evidence that the heightened sensitivity of the quadruple mutant to *Pseudomonas infection* PA14 is due to excessive oxidative stress.

## Materials and Methods

### Experimental design

The aim of this study was to determine if and how MUL-1 and its ShKT-domain paralogs regulate organismal responses to oxidative stress and DNA damage in *C. elegans*. We used genetically defined wild-type and mutant strains to compare responses to IR, chemical genotoxins, oxidative stress, osmotic stress, heat shock, starvation, and *Pseudomonas aeruginosa* infection. Stress-induced signaling and ROS levels were monitored using single copy fluorescent transcriptional and translational reporters at endogenous loci, CellROX staining, and quantitative microscopy. Strains were generated by CRISPR-Cas9 genome editing and genetic crosses. Alleles were validated by PCR and sequencing. All assays were performed with age-synchronized populations. Phenotypic outcomes were quantified using standardized assays. All experiments included at least three independent biological replicates. Sample sizes varied depending on the assay and are indicated in the corresponding figure legends. For most microscopy-based assays, 20-30 animals per condition were analyzed, whereas lifespan and progeny assays were performed using assay-appropriate cohort sizes. The number of animals (n) refers to individual worms scored per condition, unless otherwise specified.

### Strain maintenance and genetics

*Caenorhabditis elegans* strains were maintained using standard procedures as originally described by Brenner (Brenner, 1974). Animals were cultured on NGM-lite agar plates seeded with *Escherichia coli* OP50 and maintained at 20°C under standard laboratory conditions unless otherwise indicated. For specific assays, animals were propagated at 25°C (e.g., *P. aeruginosa* PA14 survival assays) or at 15°C for the maintenance of selected strains. Strains are listed under Suppl Table 2.

Age-synchronized populations were generated either by alkaline hypochlorite treatment of gravid adults followed by overnight L1 arrest in M9 buffer, or by filtration-based synchronization methods, as indicated. Transgenic and CRISPR-Cas9-edited strains were generated by standard microinjection protocols (Dokshin et al., 2018; Ghanta & Mello, 2020; Wang et al., 2018) or obtained from the *Caenorhabditis* Genetics Center (CGC) or SunyBiotech. Compound mutant strains were generated through standard genetic crosses and verified by PCR genotyping and/or DNA sequencing. All strains used in this study are listed in Suppl. Table 2, and corresponding reagents can be found in Suppl. Table 3.

### Genotoxic stress analysis

For IR experiments, age-synchronized animals were exposed to X-rays using an RS2000 X-ray irradiator (Rad Source Technologies) operated at 160 kV and 25 mA with a 0.3 mm copper filter, as previously described (Ermolaeva et al., 2013; Zou et al., 2024). Following irradiation, animals were returned to OP50-seeded NGM plates and allowed to recover under standard conditions.

For chemical genotoxic stress assays, freshly prepared aliquots of cisplatin dissolved in saline and MMS diluted in water were used according to established protocols (Volkova et al., 2020).

All solutions were protected from light until use. Groups of 20-30 age-synchronized animals were transferred into 2 mL of S-basal buffer supplemented with 5 μl of concentrated OP50 bacterial suspension as a food source. For MMS treatment, animals were exposed to 0.8 mM MMS for 16 h at 20 °C, as previously reported. For cisplatin treatment, animals were incubated with 10 μM cisplatin for 16 h at 20 °C. Samples were incubated under gentle agitation throughout the treatment period. After exposure, animals were washed thoroughly to remove residual genotoxins, transferred to OP50-seeded NGM plates, and allowed to recover.

For UV irradiation, age-synchronized animals were placed on unseeded NGM plates without lids and exposed to 200 mJ/cm^2^ UV light using a CL-1000 UV crosslinker (UVP), as previously described (Yue et al., 2024). Animals were transferred immediately to OP50-seeded NGM plates following exposure and imaged 24 h post-treatment.

### Western blot analysis of MUL-1::3xHA

Synchronized *C. elegans* populations were collected and washed three times with M9 buffer, flash-frozen in liquid nitrogen and stored at -80°C. Worm pellets were thawed on ice and mixed 1:1 with a 2x Laemmli buffer containing 5% β-mercaptoethanol, boiled at 95°C for 5 min, and briefly centrifuged. Proteins were resolved on hand-cast 12% SDS-PAGE gels in Tris-glycine-SDS buffer (Jeong et al., 2018) and transferred to a PVDF membrane using a semi-dry system (15V, ∼0.8 mA cm^-2^, 15 min). Membranes were blocked in 5% non-fat milk in PBST for 1h, incubated with mouse anti-HA (sigma, clone 16B12; 1:1000) overnight at 4°C, washed and probed with HRP-conjugated anti-mouse secondary antibody (1:5000) for 1h. Signals were detected using Pierce ECL Plus substrate and imaged on a Bio-Rad Chemidoc system.

### Microscopy and image acquisition

Images were acquired on a Zeiss Axio Imager microscope equipped with an Axiocam 503 mono camera and controlled by ZEN software. Z-stacks were collected at 1 μm intervals. For each experiment, exposure times, illumination intensity, and acquisition settings were kept constant across all genotypes and conditions. Detailed acquisition parameters for each experiment are provided in the corresponding Methods section.

### Quantification of intestinal nuclei fluorescence

Fluorescence intensity from the transcriptional reporter, and from all mutant strains generated in this reporter background, was quantified by performing line-scan measurements across the first pair of anterior intestinal nuclei in *mul-1(syb3342) IV* animals. A transverse (10 pixels wide) line was manually positioned through the nuclei, spanning 30 μm in L1-L3 larvae or 40 μm in L4 and adult animals. Line placement was optimized to minimize background contributions from adjacent intestinal cells and out-of-focus planes. For each nucleus, the maximum fluorescence intensity peak along the line profile was extracted and used for quantitative analysis.

For translational reporters and CellROX green staining, fluorescence intensity was quantified by measuring the corrected total cell fluorescence (CTCF). Images were acquired as z-stacks, and a single optical section representing a comparable focal plane was selected for each animal. Whole-animal regions of interest (ROIs) were manually delineated for individual animals, and CTCF values were calculated as the integrated fluorescence intensity after background subtraction and used for quantitative analysis.

### Stress response experiments

To assess stress responses, age-synchronized *C. elegans* at the indicated developmental stages were subjected to defined stress conditions, following established protocols.

For starvation stress, L1 larvae were transferred to unseeded NGM plates and incubated at 20 °C for 6 h, as previously described for starvation-induced stress responses (PMC3697962). For osmotic stress, synchronized L1 animals were placed on OP50-seeded NGM plates supplemented with 250 mM NaCl and incubated for 24 h at 20 °C, following standard hyperosmotic stress assays (Urso et al., 2020). For heat-shock treatment, synchronized L4 animals were incubated on OP50-seeded NGM plates at 35 °C for 1 h, followed by a 1 h recovery period at 20 °C, as previously described (Golden et al., 2020; Lithgow et al., 1995). For oxidative stress, synchronized L1 animals were exposed to 10 mM H_2_O_2_ in liquid culture. Briefly, animals suspended in M9 buffer were treated by adding 10 μl of a 5x H_2_O_2_ stock solution to 40 μl of the animal suspension and incubated for 1 h at 20 °C under gentle agitation (Offenburger & Gartner, 2018). After treatment, animals were washed four times with 1 mL M9 buffer to remove residual H_2_O_2_ transferred to OP50-seeded NGM plates, and incubated at 20 °C. Animals were imaged 24 h post-treatment.

### Generation of a stable integrated multicopy *mul-1::eGFP* line

A multicopy *mul-1::eGFP* reporter line was generated by first establishing an extrachromosomal array in the temperature-sensitive *lin-15(n765)* background using a PCR-amplified *mul-1P::linker::eGFP* fragment co-injected with the rescue plasmid pL15EK. Transgenic F1 animals showing robust intestinal GFP expression and phenotypic rescue of the *lin-15* defect were selected. To obtain a stable genomic insertion, animals carrying the array were exposed to 50 Gy of IR, and subsequent generations were screened for lines that maintained uniform GFP expression in the absence of selection, indicative of successful array integration.

### Analysis of mitochondrial ROS

A 5x CellROX green solution was prepared by diluting the 2.5 mM stock solution in M9 buffer and protected from light until use, as previously described for ROS detection in *C. elegans* (Min et al., 2021) For each reaction, synchronized L1 animals were collected in M9, and 160 μl of the suspension was mixed with 40 μl of the 5x CellROX Green solution to obtain a final reaction volume of 200 μl. Samples were incubated for 2 h at 20 °C in the dark under gentle agitation to ensure uniform staining. Following incubation, animals were pelleted by centrifugation at 1000 rpm for 1 min, washed three times with fresh M9 buffer to remove residual dye, mounted on 2% agarose pads, and imaged.

### Sensitivity to stress

Sensitivity to oxidative stress was assessed using H_2_O_2_ treatment in liquid culture following established protocols in *C. elegans* (Offenburger & Gartner, 2018). A H_2_O_2_ stock solution was freshly prepared from 30% (w/v 9.8 M) H_2_O_2_ and diluted with water to generate a 5x stock solution of 50 mM. For treatment, the 5x stock was added to synchronized L1 animals suspended in M9 buffer to obtain the desired final concentration. Samples were incubated for 1 h at 20 °C under gentle agitation. After treatment, animals were washed three times with 1 mL of M9 buffer to remove residual H_2_O_2_ and transferred to OP50-seeded NGM plates for recovery. Animals were plated in triplicate, and developmental progression was scored 48 h post-treatment.

Sensitivity to IR was assessed by exposing age-synchronized animals to X-ray irradiation, followed by analysis of post-IR developmental progression (Ermolaeva et al., 2013). Briefly, synchronized animals were irradiated with a single dose of 100 Gy, transferred to OP50-seeded NGM plates for recovery under standard conditions, and plated in triplicate. Sensitivity to IR was quantified by scoring developmental stage 48 h post-IR, using vulval morphology and overall body size as staging criteria.

### Lifespan, reproductive fitness, and apoptosis assays

Age-synchronized L4 animals from different genetic backgrounds were transferred to OP50-seeded NGM plates and maintained under standard conditions. For lifespan analysis, groups of 20 animals were plated on each NGM plate and transferred to fresh plates daily until egg-laying ceased. Viability was assessed daily by gently prodding the head or tail with a platinum wire; animals unresponsive to stimulation were scored as dead. Animals that escaped, ruptured, or died due to internal hatching (“bagging”) were censored from the analysis, following standard lifespan assay criteria (Kenyon et al., 1993).

For reproductive fitness assays, individual animals were placed on 35 mm OP50-seeded NGM plates and transferred to fresh plates daily until egg-laying ceased. After 24 h, unhatched embryos were scored as dead embryos, and after 48 h, live larvae were counted to determine brood size. Total progeny counts were analyzed and compared across genotypes, as previously described (Andux & Ellis, 2008).

For apoptosis assays, animals were collected at 24 h after the L4 stage, immobilized with 1 mM levamisole, and mounted on 2% agarose pads. Germ cell corpses in the gonad arms were visualized and quantified using fluorescence microscopy with the *ced-1::gfp(bcIs39)* reporter strain, as previously described (Zhou et al., 2001). To assess DNA damage-induced apoptosis, L4 animals were exposed to IR, and apoptotic germ cells were quantified at 24 h post-IR treatment (Gartner et al., 2000).

### *Pseudomonas* survival assays

Survival assays on *P. aeruginosa* PA14 were performed as previously described (Kwon et al., 2024) with minor modifications. Briefly, PA14 was grown overnight in LB broth at 37 °C, seeded evenly across the entire surface of NGM plates, and incubated at 37 °C for 24 h followed by an additional 24 h at 25 °C prior to use. Age-synchronized L4 animals were transferred to PA14-seeded plates supplemented with 50 μM 5-fluoro-2’-deoxyuridine (FUDR) to prevent progeny production and maintained at 25 °C. Survival was monitored every 12 h. Animals that ruptured internally (“bagging”), crawled off the agar, or exhibited vulval bursting were censored from the analysis. At least 60 animals per genotype were scored per assay, and three independent biological replicates were performed.

### Statistical analysis

Statistical analyses were performed using GraphPad Prism. For experiments involving two independent variables, data were analyzed by ordinary two-way ANOVA. When significant effects were detected, multiple comparisons were performed using Šidak’s or Tukey’s post hoc tests, as indicated in the figure legends. Comparisons between two independent groups were performed using unpaired two-tailed Student’s t-test. Survival curves were compared using log-rank (Mantel-Cox) test and the Gehan-Berslow-Wilcoxon test. All tests were two-sided. Data are presented as mean ± SEM unless stated otherwise. A p value < 0.05 was considered statistically significant.

## Acknowledgements

We want to thank the members of the Gartner Laboratory and the Korean Institute for Basic Science Center for Genomic Integrity for their fruitful discussions. We especially thank Aymeric Bailly and Albena Dinkova-Kostova for prereviewing the manuscript and Ulrike Gartner for proofreading. We thank Prof KJ Myung for his unwavering support. This work was supported by the Korean Institute for Basic Science (grant IBS-R022-D1-2025) and the National Research Foundation of Korea (grant: RS-2024-00409403). **Author contributions:** ECG, and AG: Conceptualization and writing. ECG, vast majority of reagent generation and experimental work. AGS, and KHJ, *Pseudomonas* experiments.

