## Supplementary material for "Multiple ShKT domain-containing MUL-1 proteins act as redox-responsive modulators of oxidative stress signaling in *C. elegans*": Suppl Materials

**Supplementary Materials**

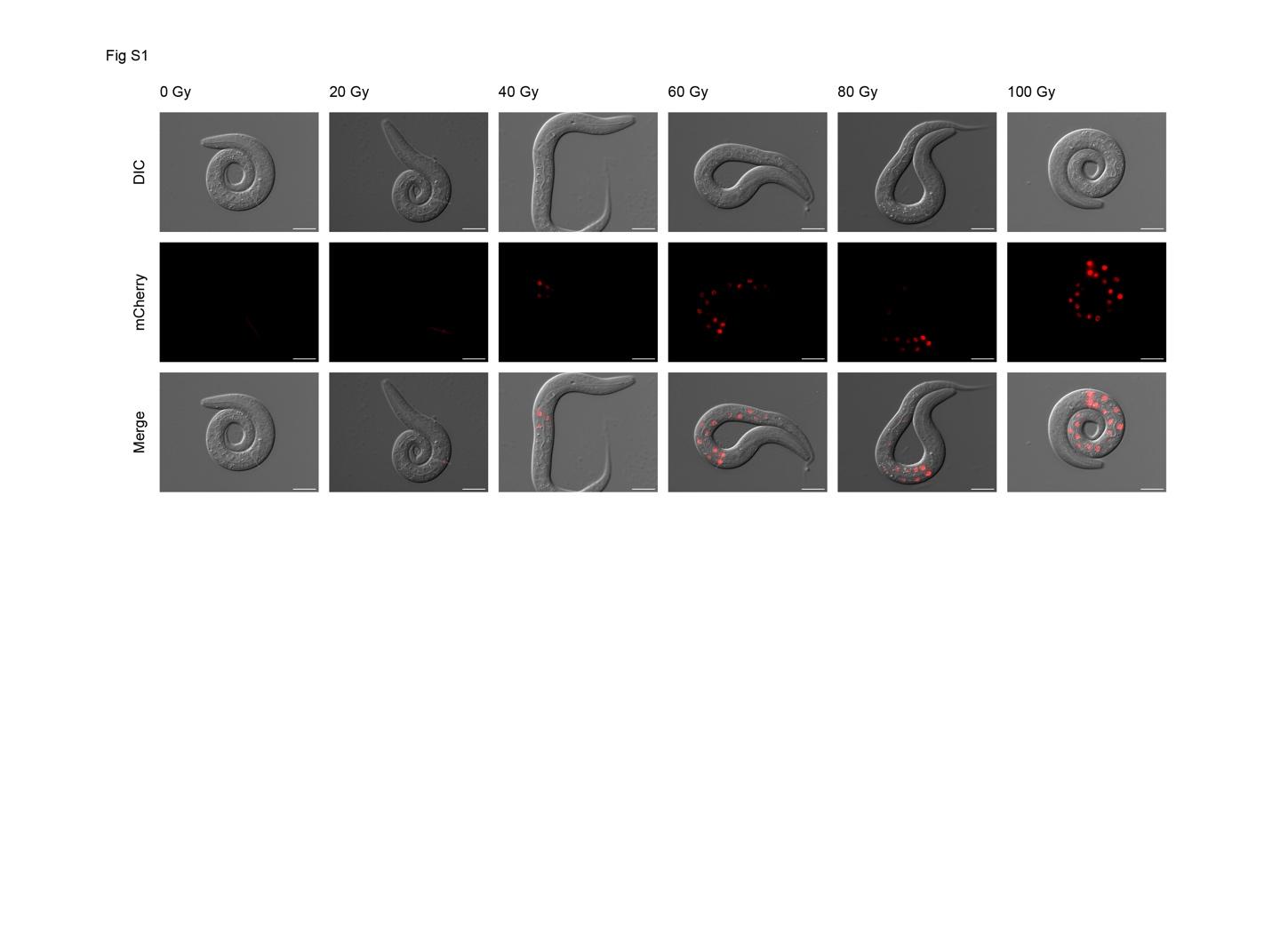

**Figure S1**

**IR induces *mul-1* expression in a dose-dependent manner.**

Representative images of *mul-1(syb3342) IV* animals 6 hours after exposure to increasing doses of IR (0-100 Gy). Reporter expression is detected in intestinal nuclei in a dose-dependent fashion, with the strongest induction observed in anterior gut cells. Representative images from three independent experiments are shown (n = 20-30 animals per condition). Scale bars, 20 µm.

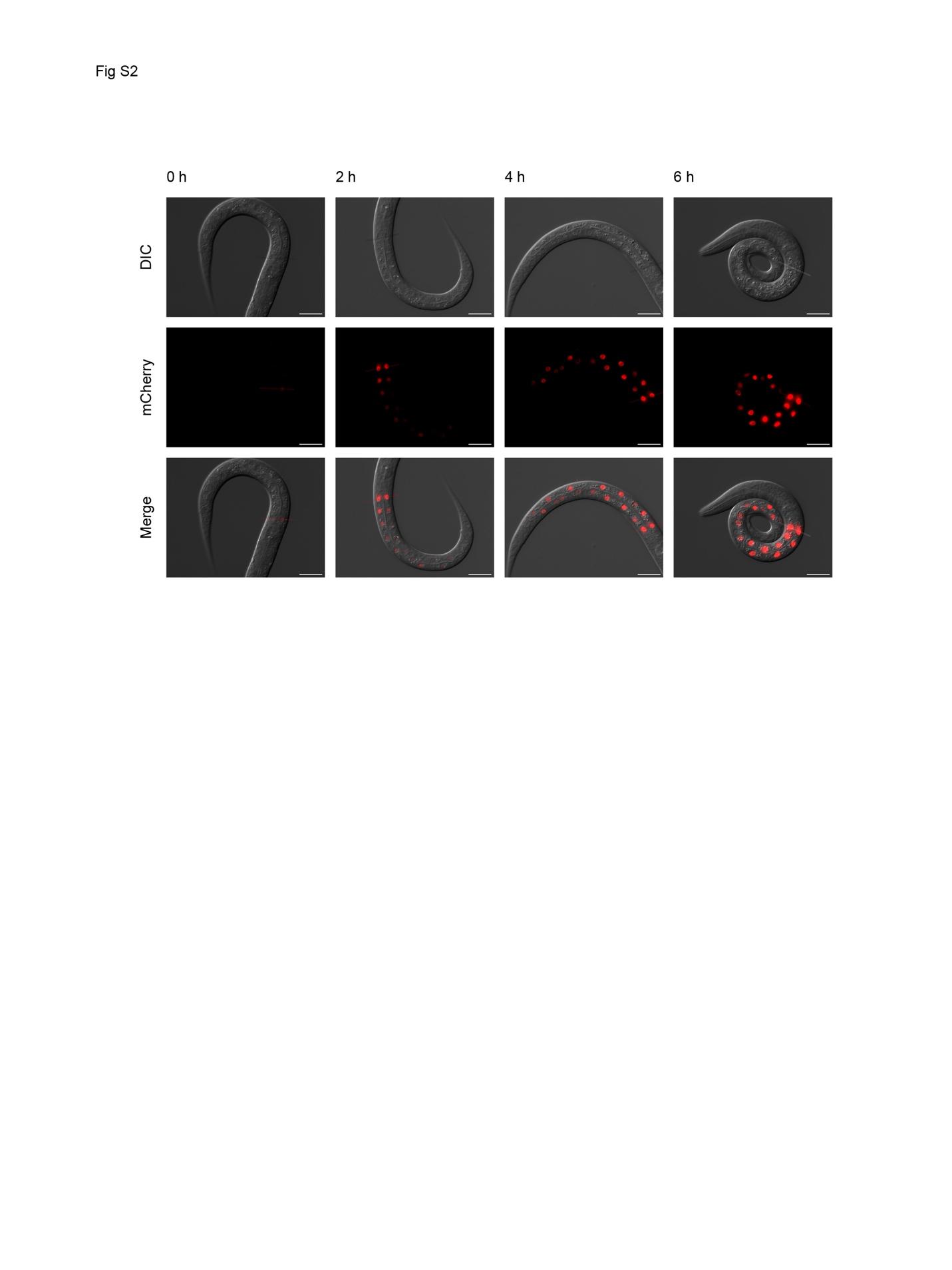

**Figure S2**

***mul-1* expression increases in a time-dependent manner following IR.**

Representative images of *mul-1(syb3342) IV* animals at indicated time points following exposure to 100 Gy IR. Reporter expression in intestinal nuclei becomes detectable at 2 hours post-IR and is strongly induced by 6 hours. Representative images from three independent experiments are shown (n = 20-30 animals per condition). Scale bars, 20 µm.

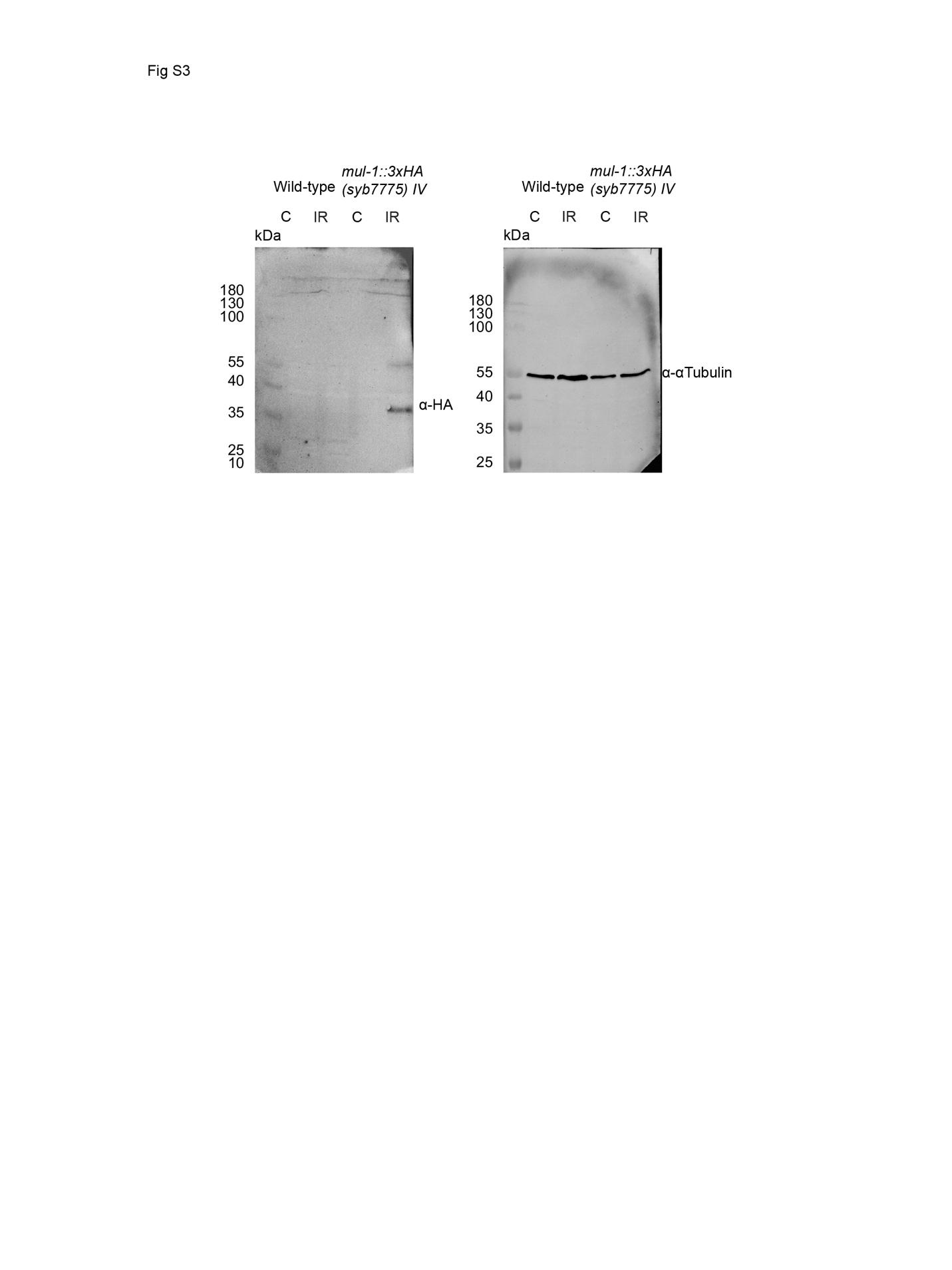

**Figure S3**

**Detection of MUL-1::3xHA protein by Western blot.**

Representative Western blot analysis of protein extracts from wild-type and *mul-1(syb7577) IV* animals under control (C) and IR conditions. MUL-1::3xHA was detected using an anti-HA antibody (α-HA), while no specific signal was observed in wild-type animals. α-Tubulin was used as a loading control. Molecular weight markers (kDa) are indicated. Representative blot from three independent experiments is shown (n = 2,000 animals per condition).

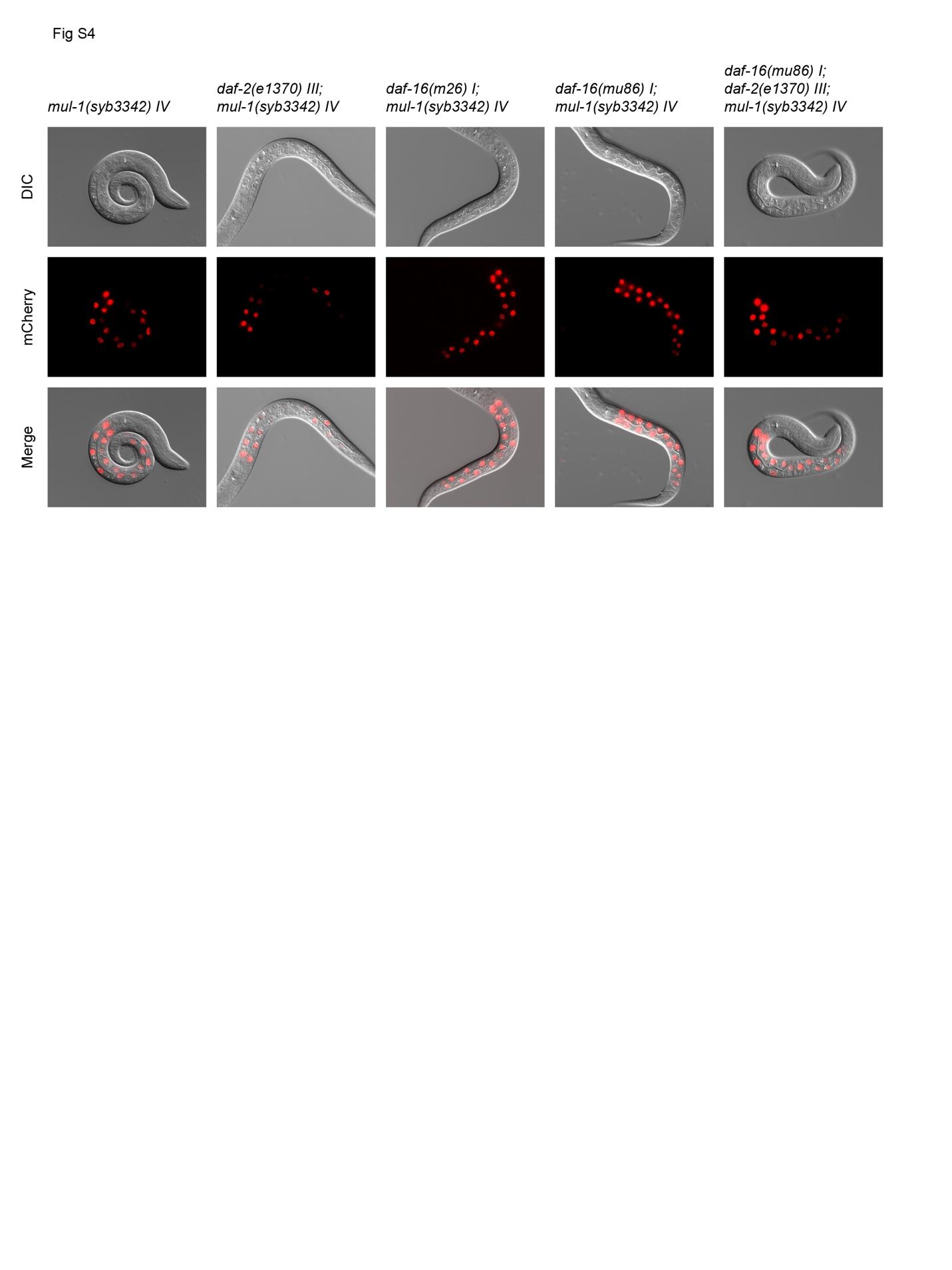

**Figure S4**

**IR-induced mul-1 expression is not dependent on the insulin/IFG signaling pathway.**

Representative images of *mul-1(syb3342) IV* animals in wild-type, *daf-2(e3170) III*, *daf-16(mu86) I*, *daf-16(m26) I*, and *daf-16(mu86) I; daf-2(e1370) III* genetic backgrounds following exposure to IR. Comparable intestinal reporter expression is observed across all genotypes, indicating that IR-dependent *mul-1* induction does not require DAF-2 or DAF-16 activity. Representative images from three independent experiments are shown (n = 20-30 animals per genotype and condition).

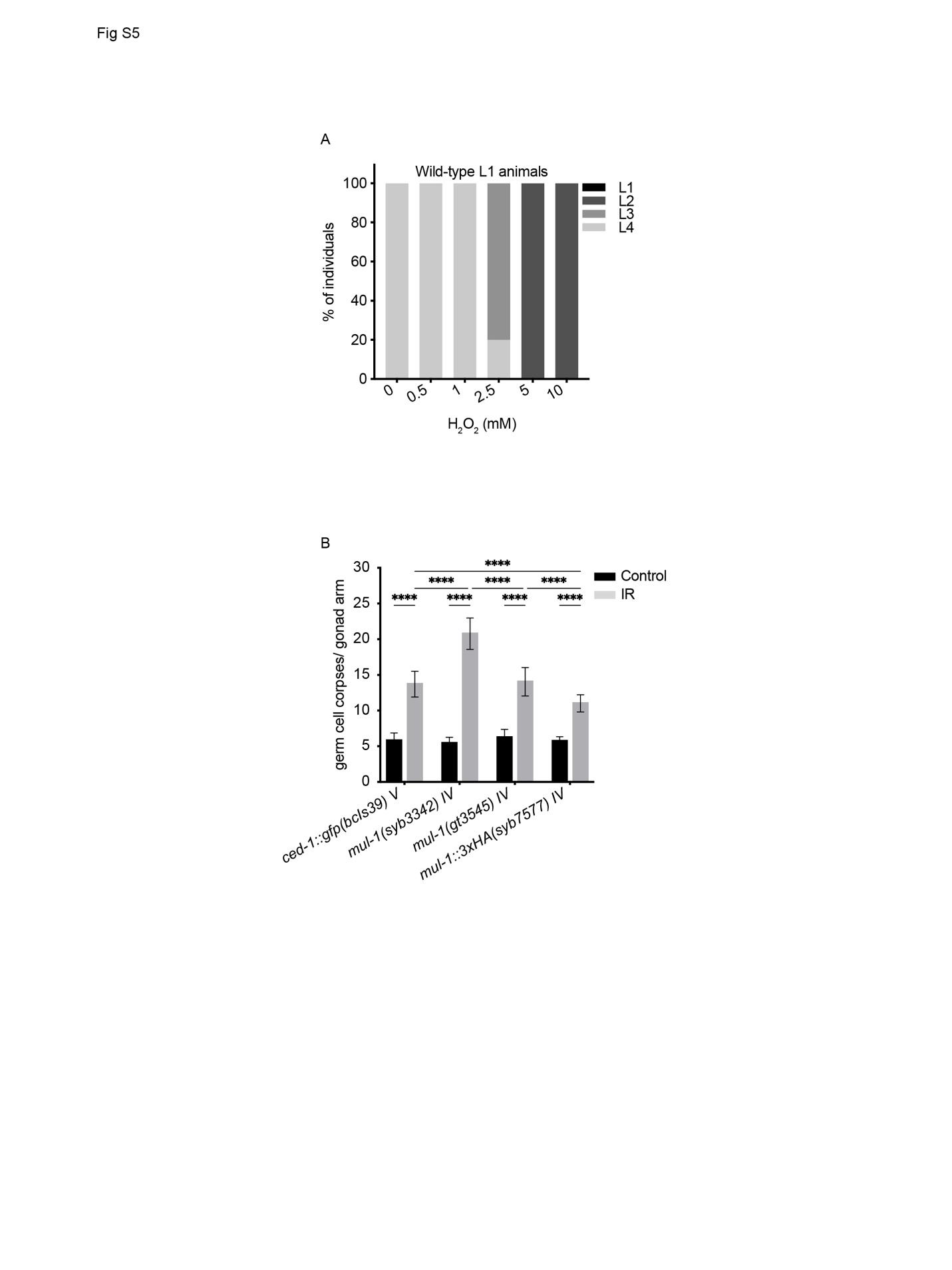

**Figure S5**

**Optimization of H_2_O_2_ dosage to assess oxidative stress sensitivity during larval development and MUL-1-tagged alleles are functionally intact.**

(A) Stage distribution of wild-type animals exposed to increasing concentrations of H_2_O_2_ at the L1 stage and scored 48 hours later. A concentration of 2.5 mM H_2_O_2_ was selected, as it induced a reproducible developmental delay without full arrest. Developmental stages were determined based on vulval morphology and body size ( n = 60 animals per condition).

(B) Quantification of germline apoptosis using the *ced-1::gfp(bcIs39)* reporter in wild-type, *mul-1(syb3342)*, *mul-1::eGFP(gt3545)*, and *mul-1::3xHA(syb7577)* animals under control conditions and following IR. Basal apoptosis levels are comparable across genotypes. Following IR, apoptosis is increased significantly in all strains; however, *mul-1(syb3342)* animals showed stronger induction of apoptotic corpses than wild-type animals. In contrast, the tagged *mul-1::eGFP* and *mul-1::3xHA* alleles did not phenocopy the elevated IR-induced apoptosis observed in *mul-1(syb3342)*, supporting the functional integrity of the tagged MUL- proteins. Data are presented as mean ± SEM from three independent experiments (n = 20-30 animals per genotype and condition). Statistical significance was determined by two-way ANOVA; ns, not significant; ****, p < 0.0001.

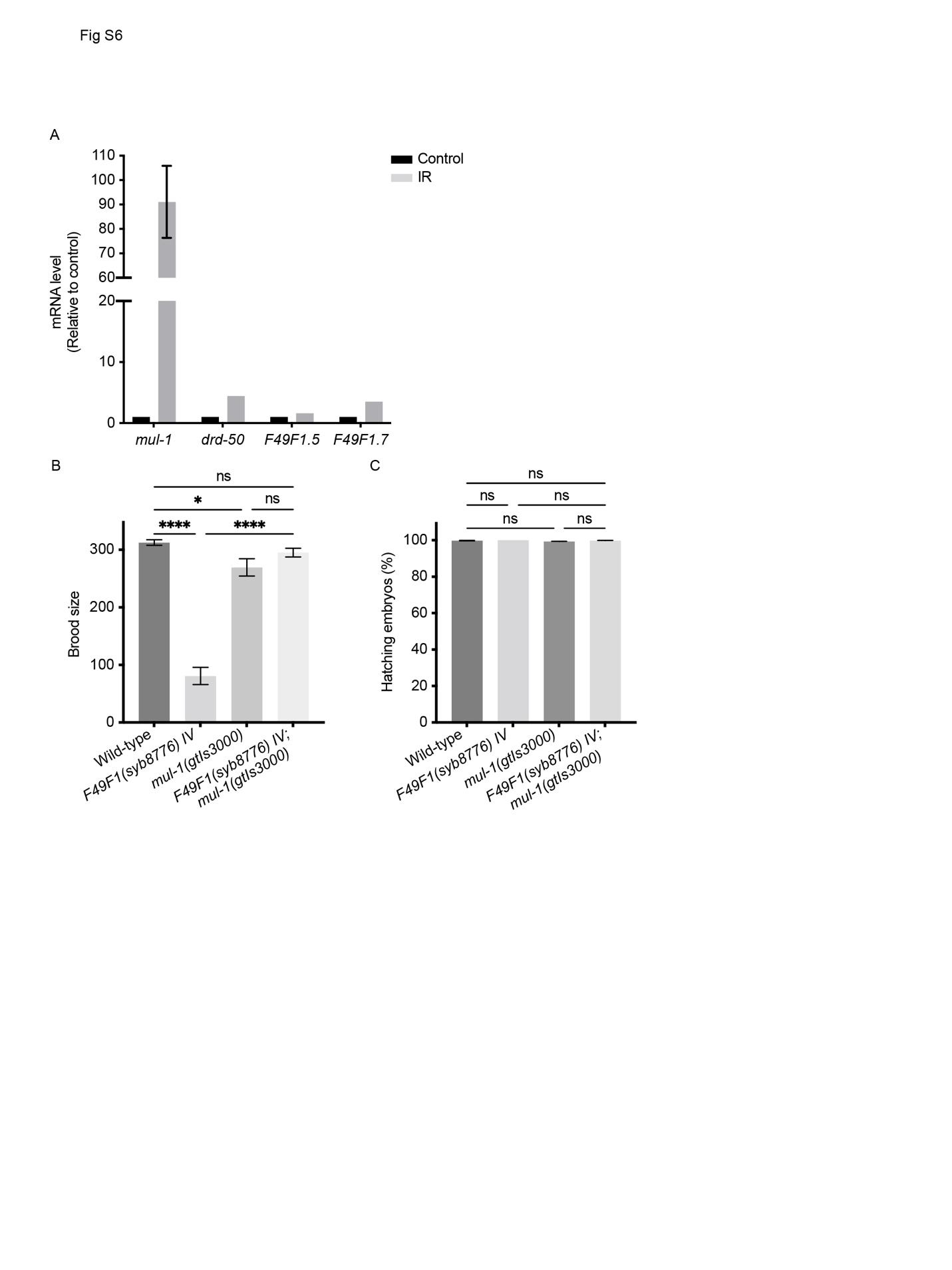

**Fig. S6**

**IR induces distinct transcriptional responses among *mul-1*-like proteins and oxidative stress markers, and multicopy *mul-1::eGFP* transgene rescues brood size of F49F1 mutants without affecting embryonic viability.**

(A) Relative mRNA expression levels of *mul-1*, its paralogs *drd-50*, *F49F1.5*, and *F49F1.7*, and the oxidative stress markers *sod-3* and *gst-4* in control and IR-treated animals. Expression levels were normalized to control conditions. IR strongly induces *mul-1* expression, whereas paralogs show only modest transcriptional changes. Data are presented as mean ± SEM from three independent experiments.

(B) Brood size analysis of wild-type, *F49F1(syb8776)*, *mul-1(gtIs3000),* and *F49F1(syb8776); mul-1(gtIs3000)* animals. The F49F1 mutant exhibits a strong reduction in progeny production compared to wild-type. Introduction of the multicopy *mul-1(gtIs3000)* array restores brood size to near wild-type levels, indicating functional rescue. *mul-1(gtIs3000)* alone does not significantly alter brood size. Data are presented as mean ± SEM from three independent experiments (n = 10-20 animals per genotype). Statistical significance was determined by one-way ANOVA with multiple comparisons; *, p < 0.05; ****, p < 0.0001; ns, not significant.

(C) Embryonic viability measured as percentage of hatching embryos in the same genotypes. No significant differences are observed across all strains, indicating that the reduced brood size in F49F1 mutants is not due to increased embryonic lethality. Data are presented as mean ± SEM from three independent experiments (n = 10-20 animals per genotype).

**Suppl. Table 1**

**MUL-1 paralogs**

| **Accession** | **Name** | **Locus** | **Genomic location** | **ID Rank** | **E value** |
| --- | --- | --- | --- | --- | --- |
| NP_500487.2 | ***mul-1a*** | **F49F1.6a** | IV:4121342..4123166 | 0 | 0 |
| NP_001299956.1 | ***mul-1b*** | **F49F1.6b** | IV:4121342..4123166 | 1 | 9E-133 |
| NP_500485.1 | ***drd-50*** | **F49F1.1** | IV:4117732..4119179 | 2 | 6E-60 |
| NP_500488.1 |  | **F49F1.7a** | IV:4123525..4125757 | 3 | 3E-37 |
| NP_507973.1 |  | **F46B3.1** | V:20596210..20597176 | 4 | 3E-26 |
| NP_500486.1 |  | F49F1.5a | IV:4119460..4120415 | 5 | 6E-16 |
| NP_504146.1 |  | T05B4.9 | V:4117585..4118432 | 6 | 2E-13 |
| NP_496932.1 |  | F01D5.1 | II:13996958..13997581 | 7 | 1E-13 |
| NP_504145.2 |  | T05B4.8 | V:4115089..4116171 | 8 | 4E-12 |
| NP_504147.1 |  | T05B4.10 | V:4119890..4120729 | 9 | 5E-12 |
| NP_001294421.1 |  | F49F1.5b | IV:4119460..4120415 | 10 | 2E-12 |
| NP_496930.1 |  | Y39G8B.7 | II:13994410..13995034 | 11 | 8E-12 |
| NP_507329.1 |  | ZK218.1 | V:17092562..17093546 | 12 | 4E-11 |
| NP_507331.1 |  | ZK218.3 | V:17097800..17098769 | 13 | 6E-11 |
| NP_001022431.2 |  | Y39G8B.10 | II:13982048..13983115 | 14 | 1E-11 |
| NP_504149.1 |  | T05B4.12 | V:4123171..4126034 | 15 | 8E-11 |
| NP_001254248.1 |  | ZK673.1a | II:10443146..10445415 | 16 | 3E-11 |
| NP_504148.1 | *phat-5* | T05B4.11 | V:4121489..4122515 | 17 | 3E-10 |
| NP_504978.1 |  | F07C4.10 | V:7638894..7639887 | 18 | 6E-10 |
| NP_506977.2 |  | F35E8.1 | V:15904192..15905630 | 19 | 6E-10 |
| NP_504877.2 | *tag-293* | C03G6.13 | V:7354435..7355564 | 20 | 6E-10 |
| NP_507333.1 |  | ZK218.5 | V:17101129..17102109 | 21 | 0.000000001 |
| NP_001254246.1 |  | E04D5.4a | II:10440276..10442517 | 22 | 6E-10 |
| NP_507335.1 |  | ZK218.7 | V:17105783..17106656 | 23 | 0.000000002 |
| NP_507336.2 |  | ZK218.11 | V:17107388..17108410 | 24 | 0.000000002 |
| NP_504979.2 |  | F07C4.11 | V:7642519..7643516 | 25 | 0.000000003 |
| NP_001024810.2 |  | M163.11 | X:14479378..14480256 | 26 | 0.000000004 |
| NP_001368262.1 |  | F49F1.7b | IV:4123525..4125757 | 27 | 0.000000002 |
| NP_506985.1 |  | F35E8.10 | V:15919434..15920373 | 28 | 0.000000007 |
| NP_506976.1 |  | T06C12.14 | V:15899996..15901054 | 29 | 0.00000001 |
| NP_001317750.1 |  | T05B4.13 | V:4126319..4127535 | 30 | 0.00000001 |
| NP_506984.1 |  | F35E8.9 | V:15916334..15917558 | 31 | 0.00000002 |
| NP_001379423.1 |  | M163.8 | X:14485127..14485955 | 32 | 0.000000004 |
| NP_504129.1 | *phat-3* | C49G7.4 | V:4043932..4044951 | 33 | 0.00000003 |
| NP_504130.1 | *phat-7* | C49G7.3 | V:4046422..4047433 | 34 | 0.00000004 |
| NP_001123161.2 |  | K03A11.6 | X:13055860..13058381 | 35 | 0.0000004 |
| NP_001023993.1 | *nas-31* | F58B4.1a | V:10918857..10921847 | 36 | 0.0000008 |
| NP_496935.1 |  | F01D5.5 | II:14001686..14002300 | 37 | 0.0000003 |
| NP_001022432.1 |  | Y39G8B.9 | II:13981365..13981785 | 38 | 0.0000001 |
| NP_506981.1 |  | F35E8.6 | V:15912148..15913056 | 39 | 0.000001 |
| NP_001254149.1 |  | M05D6.8b | II:8470076..8471239 | 40 | 0.0000006 |
| NP_495937.2 | ***sysm-1*** | T24B8.5 | II:9082626..9083193 | 41 | 0.000003 |
| NP_503462.1 |  | Y46H3D.8 | V:1630309..1631248 | 42 | 0.00004 |
| NP_496933.1 |  | F01D5.2 | II:13998460..13999064 | 43 | 0.00002 |
| NP_001255258.1 |  | K08D10.13 | IV:4170728..4171581 | 44 | 0.00002 |
| NP_510745.2 |  | F52G3.5 | X:16973226..16976143 | 45 | 0.0003 |
| NP_496934.1 |  | F01D5.3 | II:13999059..14000144 | 46 | 0.0001 |
| NP_501871.2 | *nas-12* | C24F3.3 | IV:10224023..10226387 | 47 | 0.0008 |
| NP_504152.1 | *phat-4* | T05B4.3 | V:4129904..4130909 | 48 | 0.0006 |
| NP_499332.2 |  | Y45F3A.8 | III:10589865..10592829 | 49 | 0.003 |
| NP_491507.2 |  | C46H11.7 | I:5042102..5043064 | 50 | 0.003 |
| NP_507155.1 |  | F26D2.13 | V:16434734..16435610 | 51 | 0.004 |
| NP_507156.2 |  | F26D2.14 | V:16435855..16436736 | 52 | 0.008 |
| NP_498981.2 |  | ZK643.6 | III:8957701..8960764 | 53 | 0.009 |
| NP_507558.1 |  | F16H6.3 | V:18195306..18197863 | 54 | 0.043 |

**Suppl. Table 2**

**List of strains**

| REAGENT or RESOURCE | SOURCE | IDENTIFIER |
| --- | --- | --- |
| Experimental models: Organisms/strains |  |  |
| *C. elegans*: Wild-type N2 | *Caenorhabditis* Genetics Center (CGC) | WormBase ID: WBStrain00000001 |
| *Escherichia coli* OP50 | CGC |  |
| *Pseudomonas aeruginosa* PA14 | Gift from Seung Jae V. Lee |  |
| *mul-1(syb3342 [mul-1P::mCherry::linker::H2B]) IV* | This study | PHX3342 |
| *mul-1(syb7577[mul-1::3xHA]) IV* | This study | PHX7577 |
| *prdx-2(gk169) II; mul-1(syb3342) IV* | This study | TG5005 |
| *mul-1(syb3342) IV, sek-1(km4) X* | This study | TG5001 |
| *mul-1(syb3342) pmk-1(km25) IV* | This study | TG4689 |
| *mul-1(syb3342) skn-1(zj15) IV* | This study | TG4881 |
| *atf-7(q22q130) III; mul-1(syb3342) IV* | This study | TG4705 |
| *daf-2(e1370) III; mul-1(syb3342) IV* | This study | TG4877 |
| *daf-16(m26) I; mul-1(syb3342) IV* | This study | TG4878 |
| *daf-16(mu86) I; mul-1(syb3342) IV* | This study | TG4942 |
| *daf-16(mu86) I; daf-2(e1370) III; mul-1(syb3342) IV* | This study | TG4989 |
| *glo-1(gt3565) X* | This study | TG5064 |
| *mul-1(gt3545 [mul-1::linker::eGFP) IV; glo-1(zu391) X* | This study | TG5065 |
| *mul-1(gtIs3000[mul-1P::mul-1::eGFP]) lin-15(+)* | This study | TG5231 |
| *prdx-2(gk169) II; mul-1(gt3545) IV; glo-1(zu391) X* | This study | TG5066 |
| *daf-16(mu86) I* | CGC | CF1038 |
| *pmk-1(km25) IV* | CGC | KU25 |
| *bcIs39 [lim-7P::ced-1::GFP + lin-15(+)] V* | CGC | MD701 |
| *cep-1(lg12501) I* | CGC | XY1054 |
| *cep-1(lg12501) I; bcIs39 V* | This study | TG5286 |
| *mul-1(syb3342) IV; bcIs39 V* | This study | TG4611 |
| *mul-1(gt3457[mul-1::STOP-IN]) IV* | This study | TG4910 |
| *mul-1(gt3459[mul-1::STOP-IN]) IV; bcIs39 V* | This study | TG4912 |
| *cep-1(lg12501) I; mul-1(syb3342) IV; bcIs39 V* | This study | TG4927 |
| *F49F1(syb8776) IV* | This study | PHX8776 |
| *F49F1(syb8776) IV; bcIs39 V* | This study | TG5122 |
| *cep-1(lg12501) I; F49F1(syb8776) IV; bcIs39 V* | This study | TG5256 |
| *gst-4(gt3596[gst-4P::gst-4::mCherry]) IV; sod-3(gt3598[sod-3P::sod-3::eGFP]) glo-1(zu391) X* | This study | TG5166 |
| *mul-1(gt3625[mul-1::STOP-IN]) gst-4(gt3596) IV; sod-3(gt3598) glo-1(zu391) X* | This study | TG5183 |
| *F49F1(gt3613) gst-4(gt3596) IV; sod-3(gt3598) glo-1(zu391) X* | This study | TG5171 |
| *prdx-2(gk169) II; gst-4(gt3596) IV; sod-3(gt3598) glo-1(zu391) X* | This study | TG5175 |
| *prdx-2(gk169) II; skn-1(zj15) gst-4(gt3596) IV; sod-3(gt3598) glo-1(zu391) X* | This study | TG5234 |
| *daf-16(mu86) I; prdx-2(gk169) II; gst-4(gt3596) IV; sod-3(gt3598) glo-1(zu391) X* | This study | TG5233 |
| *sysm-1(ok3236) II; bcIs39 V* | This study | TG4923 |
| *sysm-1(ok3236) II; mul-1(syb3342) IV; bcIs39 V* | This study | TG5245 |
| *sysm-1(ok3236) II; F49F1(syb8776) IV; bcIs39 V* | This study | TG5242 |
| *F49F19syb8776) IV; F46B3.1(syb8669) V* | This study | TG5121 |

**Suppl. Table Table 3**

**List of oligonucleotides and reagents**

| Oligonucleotide | Sequence | Application |
| --- | --- | --- |
| AG24-seq-s | TCACAAGTTCGACCCCCAGT | Genotyping |
| AG24-seq-a | CTGCGTCTAAATGCCTGCAAT | Genotyping |
| mCherry-mid-s | GACGGAGGAGTTGTTACAGTGAC | Genotyping |
| prdx-2 L crRNA | agcgacgaaagaaaaaaaat | Genome editing |
| prdx-2 R crRNA | TAGCCTTGAACTCCTCAGCA | Genome editing |
| prdx-(gk169) ssODN | cccagaatgttttattagttctcatttcgtcctccgattttttttctttcAACACCGTTGTGCTCGCCGCTTCCACCGACTCTGTCTTCTCTCACTTGGC | Genome editing |
| prdx-2 F | gaaatcatgtctctcgctcc | Genotyping |
| prdx-2 Int F | TGACTTCACTTTCGTGTGCC | Genotyping |
| prdx-2 R | ATTTGGTGGTTGGTGTCAGC | Genome editing |
| sek-1 crRNA L | atacactagaataagtgact | Genome editing |
| sek-1 crRNA R | gcatagtttttacctaacta | Genome editing |
| sek-1(km4) ssODN | agatttatcaatacttactgatttcaaaaattcattttcatccatacactagaataagtgctatgctagatttgcgaaaaattgcaaaaacttctgaattcattcattgc | Genome editing |
| sek-1 F | ttcctatttggctggtgcac | Genotyping |
| sek-1 R | GCCAAACAGTGTCGAGAATc | Genotyping |
| sek-1 Int F | agtttgtttccagcaggacc | Genotyping |
| pmk-1 crRNA L | acaaacatgattccaggtaa | Genome editing |
| pmk-1 crRNA R | GGAGTTCACGATATGTACGA | Genome editing |
| pmk-1 ssODN | catttaatcggttcaaattgccgcttaatttttatatttacatttatattggttttacgtCGTTGTATGTGTCATGAAAATATAATTGATCTACTTGATG | Genome editing |
| pmk-1 F | aagttgccatgacctcagag | Genotyping |
| pmk-1 R | GTCTGCGTGTAATGCATCCA | Genotyping |
| skn-1 crRNA | TTATAATCAGGCAAATTgta | Genome editing |
| skn-1(zj15) ssODN | TCCAGTTATCAATAATGTTTCTCTGTCGGAAGGAATTGTTTATAATCAGGCTAATTgcatggttttgatttaattattgtgttttcattcaacattttttatttttcagT | Genome editing |
| skn-1 F | ACTCACTGCCGAAGAGAATG | Genotyping |
| skn-1 R | agaggagaaagtgcaaccag | Genotyping |
| atf-7 crRNA | ACATCGTTGCATTAGTTCAG | Genome editing |
| atf-7(qd22qd130) ssODN | TCCAAACCCAAAATTAATGTTCACTCCACTTGATCTACCAACAACCGCTGAGCTAATGCAGCGATGTCTAGCAGTAAATTCATTCGAAGCAAAGTTCTGAGAAGCGAATCAAAAG | Genome editing |
| atf-7 F | gccaacgtcgatgacaaatg | Genotyping |
| atf-7 R | GAAATGTCCGCGGTTTTCAG | Genotyping |
| daf-2 crRNA | TATTGGTTTGAGTAATGATG | Genome editing |
| daf-2(e1370) ssODN | TGTTCTCTATGAAATGGTTACACTCGGTGCTCAGTCATATATTGGTTTGAGTAACGATGAAGTGTTGAATTATATTGGAATGGCCCGGAAGGTTATCAAGAAGCCCGAAT | Genome editing |
| daf-2 F | GACTATCTCCGATCGAAACG | Genotyping |
| daf-2 R | GAATGACTCGTCGTCGCATT | Genotyping |
| daf-16 crRNA | TCATCAGACATCGTTTCCTT | Genome editing |
| daf-16(m26) ssODN | ACAGCAATGCTTCATACTCCAGATGGAAGCAATTCTCATCAGACATCCTTTCCTTCTGAatgagctttttcataattattttttggagatttaattaatattgaaacatt | Genome editing |
| daf-16 F | CTATACGGGAGCAATGAGCA | Genotyping |
| daf-16 R | GGTCGTTGTCTTTTTTCCCG | Genotyping |
| daf-16 crRNA L | ctagtcggactgacgtagat | Genome editing |
| daf-16 crRNA R | CAAAGCCAGGAAGGAATCCA | Genome editing |
| daf-16 (mu86) ssODN | ctgtgtgccttccttttttgattccaatctcaaagattttcccacattcgtgtgggttttctagtcggacCACGGCGTACACGTGAACGATCCAATACTATTGAGACGACTACAAAGgtaagagatagtgaaaataatttt | Genome editing |
| daf-16 mu86 F | tagacggtgaccatctagag | Genotyping |
| daf-16 mu86 R | CCAATAGCTGGAGAAACACG | Genotyping |
| daf-16 mu86 Int F | atcttgcagAACTCGATCCG | Genotyping |
| glo-1 crRNA | ATATTGACCACCTTGTTCAG | Genome editing |
| glo-1(zu391) ssODN | CGTATTTTCCCTTGAAGACCTTTTTTTAGTCATTTAAATTACTAATGTTTTCAAGTGATCTCCACCGAACAAGGTGGCCAATATGATGTCCCTTTCATGAATCGTGAAGGCAACGTGAATCTTGATGACAATACCAC | Genome editing |
| glo-1 zu391 F | gtcctcaacgatgtaatcac | Genotyping |
| glo-1 zu391 R | gttcacggtcatTTAGCAAC | Genotyping |
| mul-1 crRNA 3 | CTATCCACCGGAGAAGAGAA | Genome editing |
| mul-1 linker eGFP F | GCCGGGTGGGCAAAAAACGGCTTCTGCACGAACACCTTCTATCCGCCAGAGAAAAGGAAGGAGTACTGTGCAAAGACATGTAGAATGTGCGGAGGTGGAGGTTCCGGTGGAGGTGGTTCCTCCAAGGGAGAGGAGCTCTT | Genome editing |
| eGFP mul-1 donor 120bp R | aagaagtcataattcctcatgttcgacttcacaccctcttttgtcatcacattgatacacattttgataagaataaataatttattaaatcaaaaaaTTACTTGTAGAGCTCGTCCATTC | Genome editing |
| EGFP 1-2-3 F | CCAAGGGAGAGGAGCTCTTCA | Genome editing |
| EGFP 1-2-3 R | CTTGTAGAGCTCGTCCATTC | Genome editing |
| mul-1 gen F | GCAAGGATTCGTCGCCAAAg | Genotyping |
| mul-1 gen R | cggacaaatacattcacacg | Genotyping |
| EGFP Internal F | CATGCCAGAGGGATACGTCC | Genotyping |
| EGFP Internal R | GGACGTATCCCTCTGGCATG | Genotyping |
| mul-1 Nest F | GCGAAGAATGGCTTTTGCAC | Genotyping |
| mul-1 Nest R | CAGAAGGTTCAAAGCAAGCG | Genotyping |
| mul-1 multi F | gatttttctgcgataacttcag | Genotyping |
| mul-1 multi R | gaattgataatagtgtctgtctgc | Genotyping |
| mul-1 060722 F | TGGGCAAAAAACGGCTTCTG | Genotyping |
| mul-1 060722 R | cgacttcacaccctcttttg | Genotyping |
| cep-1 DK F | AATGTATCCAGGCGCAGTTC | Genotyping |
| cep-1 R | ACAGCGACTTCTCTTCATCG | Genotyping |
| cep-1 DK R | aacatcattcgtagccggac | Genotyping |
| F49F1 crRNA L | TTTGTACCATTTGTGACTTG | Genome editing |
| F49F1 crRNA R | AGATAAAAATAATGAAAAGC | Genome editing |
| F49F1 IV ssODN | TAACGCTGCCTTTTTATAGAACTTTGATTATCAGAAAATAGTAGCTCTTTGAACATTTTCCAATGATTTGAGAATTTTGGAGCAAAGTTTTGAAGGTTTTCTTATCAGTA | Genome editing |
| F49F1 F | TCACATCATCATTTCCCCGC | Genotyping |
| F49F1 Int R | TTTAATTGGGCAGAGGCTGG | Genotyping |
| F49F1 R | GAGCTGTATTCGGAGAAGAG | Genotyping |
| mul-1 2 crRNA | agACAATGCCCATACACCTG | Genome editing |
| mul-1 STOP IN ssODN | atgagcattcataaatcaaactacctttatttccagACAAGGGAAGTTTGTCCAGAGCAGAGGTGACTAAGTGATAAGCTAGCTGCCCATACACCTGTGGAACCTGCGTACCAGCTACCC | Genome editing |
| mul-1 F qPCR | CTTGTGGAATATGCCATCAG | Genotyping |
| mul-1 11072022 R | CGTCAGTACATGGAACGCTA | Genotyping |
| F46B1.3 F | gaccactcctgtgtttgaac | Genotyping |
| F46B1.3 Int F | AAAGTGTCCACTGAGCTGTG | Genotyping |
| F46B1.3 R | ttgtggctgtgagttgaagc | Genotyping |
| gst-4 crRNA R | GAATTACGTGGCAACAAGAA | Genome editing |
| gst-4 120 bp F | AGATTTCAACAAAGAAAAGAAGTTGGAAGAGTTCTATAACAAGATTCATTCAATTCCAGAAATTAAGAATTACGTAGCAACGAGAAAAGATAGTATTGTTGTCTCAAAGGGTGAAGAAGA | Genome editing |
| gst-4 120 bp R | ataataataatttattacgctctctgggagacgtgataggaacgatattattttactacatacataattcagacttaaataattcgattTTACTTATACAATTCATCCATGC | Genome editing |
| gst-4 10192023 F | AACCAGCCCGTGATGATTTC | Genotyping |
| mCherry F | GTCTCAAAGGGTGAAGAAGA | Genome editing |
| mCherry R | CTTATACAATTCATCCATGC | Genome editing |
| sod-3 crRNA | CAATGCTCGACAATAAaagc | Genome editing |
| sod-3 120 bp F | GgtgatttttgagttttgaaagcgtcgatttttaatataaaaaaattttcagATTGCCAACTGGAAGAATATCAGCGAGAGATTTGCCAATGCTCGACAATCCAAGgtaacacttagttt | Genome editing |
| sod-3 120 bp R | taaacaattacaaaaatacaaattctcagcgttttaaactacatctgaattatcttcaatattttcgtaaaaccgaaaattccaatatttcctgcttTTACTTGTAGAGCTCGTCCATTC | Genome editing |
| sod-3 F | TGGGGATGGTTGGGATATTG | Genotyping |
| sod-3 R | cagtgtaccgagtgaagttc | Genotyping |
| eGFP Int 10192023 F | TCAACCGTATCGAGCTCAAG | Genotyping |
| Reagent | Supplier | Catalog Number |
| cis-Diamminedichloroplatinum(II) dichloride (Cisplatin) | Sigma-Aldrich | P4394-250MG |
| Methyl methanesulfonate (MMS), 99% | Aldrich, Sigma-Aldrich | 129925-25G |
| Hydrogen peroxide 30% | Supelco/ Merck, Sigma-Aldrich | 1.07209.0250 |
| Tetramisole hydrochloride ≥99% | Sigma | L9756-6G |
| KOD FX Neo DNA Polymerase | TOYOBO | KFX-201 |
| CellROX™ Green Reagent (2.5 mM in DMSO) | Invitrogen, Thermo Fisher | C10444 |
